# Proteome-wide computational analyses reveal links between protein condensate formation and RNA biology

**DOI:** 10.1101/2025.03.03.640993

**Authors:** Snigdha Maiti, Swarnendu Tripathi, David W Baggett, Aaron H. Phillips, Cheon-Gil Park, Jina Wang, Wahiduzzaman, William T Freyaldenhoven, Swati Kinger, Brittany Pioso, John Bollinger, Ramiz Somjee, Benjamin Lang, M. Madan Babu, Richard W. Kriwacki

## Abstract

Biomolecular condensates mediate dynamic compartmentalization of cellular processes. The multivalent interactions that underlie biomolecular condensation are often promoted by intrinsically disordered regions (IDRs) within proteins. While the role of IDRs in biomolecular condensates is well appreciated, predicting whether an IDR forms condensates in cells remains challenging. Here, we developed a machine learning model to predict condensation behavior of IDRs, analyzing 215 IDRs from fusion oncoproteins in HEK293T cells. Our study identified distinct sequence-derived physicochemical features associated with condensation. Leveraging these data, our model predicts that ∼12% of the ∼13,000 IDRs in the human proteome are likely to form cellular condensates. Proteins with condensate-forming IDRs are enriched in functions involving RNA-related processes and membrane-less organelles (MLOs), highlighting their role in MLO assembly and function. Our model, available via the SAK3.0 web server (https://sak.stjude.org), provides a powerful resource for studying IDR-driven phase separation across proteomes, offering insights into biomolecular condensates and their biological roles.

## Introduction

Biomolecular condensates, also termed membrane-less organelles (MLOs), mediate dynamic compartmentalization of biomolecules associated with diverse biological processes^1^. Found in both the nucleus (nucleoli, paraspeckles, Cajal bodies, PML bodies) and the cytoplasm (P-bodies, stress granules, germ granules), MLOs are essential for myriad cellular functions, and their dysfunction has been implicated in numerous diseases^2–5^. Assembly of biomolecules into condensates is often driven by phase separation (PS), which is facilitated by weak, multivalent interactions between constituent biomolecules^6,7^ and often promoted by intrinsically disordered regions (IDRs) in proteins^8^. IDRs lack stable secondary and tertiary structures and often retain a high degree of disorder when interacting with other molecules to form condensates^9–11^. They show biased amino acid composition, with depletion in structure-promoting hydrophobic amino acids and enrichment of polar and charged amino acids, features that enable the dynamic multivalent interactions that underpin phase separation^12^. Analyses of amino acid patterns in IDRs across diverse biomolecular condensates have revealed the sequence-related physicochemical features that govern condensate formation^13–16^.

Multiple algorithms have been developed to identify those features in IDRs associated with PS and several predictors are available^17–27^. For example, PhaSePred^22^ predicts PS in autonomously self-assembling and partner-dependent proteins using multimodal features. DeePhase^23^ utilizes a combined engineered features model and a neural network-based language model to predict the propensity of proteins to undergo homotypic PS. FuzDrop^28^, a conformational entropy-based model, predicts droplet promoting regions and proteins driving PS. Tools including PScore^17^, ParSe v2^25^, and catGranule^29^, leverage sequence-specific features, such as sequence composition, prion-like domains, and charged residues, while others such as MolPhase^30^ integrate multidimensional data, including structural and physicochemical properties. Notably, LLPhyScore^21^ predicts protein PS potential using a comprehensive scoring system based on sequence and physicochemical features of IDRs, offering valuable insights into mechanisms of IDR-driven protein phase separation. A recently developed predictor called PICNIC^31^ utilizes protein-protein interaction networks to exclude proteins that have a connection with known condensate-forming proteins. However, these predictors rely on literature-mined data, which were often obtained using non-standard experimental conditions, leading to variability and noise. Additionally, these studies often use information from the Protein Data Bank (PDB)^32^ or the human proteome as negative datasets, which results in false negatives. Curating a verified set of IDRs, including those that form condensates and others that remain diffuse under controlled experimental conditions, is crucial for training and validating predictive models^33^. In our previous study, using experimentally validated datasets, we developed the FO-Puncta ML model to predict the condensation behavior of fusion oncoproteins (FO) in cells^34^. However, this model does not identify the specific regions responsible for condensate formation, nor does it elucidate the role and influence of IDRs present in these FOs on the condensation process. Recently, a machine learning model was developed that predicts homotypic PS of IDRs^35^, but heterotypic interactions (between IDRs and other proteins or nucleic acids) contribute to the formation of many biological condensates. Thus, it is important to assess IDR condensate formation under conditions that reflect the compositional complexity of cellular environments.

We addressed these challenges by examining the cellular condensation behavior of 215 IDRs derived from 149 human fusion oncoproteins (FO), 58% (96) of which were previously shown to form cellular condensates^34^. Each IDR was tagged with monomeric, enhanced green fluorescent protein (GFP) and assessed in cells using standardized conditions and microscopy protocols, coupled with rigorous computational image analysis. These efforts yielded a dataset of condensate-forming [termed puncta(+)] and condensate-negative [puncta(-)] IDRs, and the data was used to develop a machine learning (ML) model (termed IDR-Puncta ML model) to predict IDR condensation with high accuracy, based on a large set of sequence-derived physicochemical features. Using the IDR-Puncta ML model, we demonstrate that 12% of 12,899 IDRs identified in the human proteome (termed the human “IDRome”) are predicted to form biomolecular condensates. We show that proteins containing puncta(+) IDRs (1,393 of 8,067 IDR-containing human proteins) are significantly enriched in Gene Ontology (GO) terms involving RNA-related biological processes, including transcription and processing, and others associated with cell division and actin cytoskeleton, and are over-represented as constituents of MLOs. These findings indicate that proteins with puncta(+) IDRs are specialized for functions associated with nuclear processes involving RNA processing, suggesting a role for PS-driven compartmentalization of proteins, RNA, and other biomolecules in these processes. We note that these findings are based on analyses of IDRs derived from cancer-associated FOs and their functional associations may be biased toward biological processes altered in cancer. However, the physicochemical features of FO-derived IDRs broadly sample those of the human IDRome, supporting the generality of our conclusions on the specialized biological functions of puncta(+) IDR-containing human proteins. Finally, the IDR-Puncta ML model, given its high accuracy, will be a powerful tool for uncovering the biological roles of proteins with puncta(+) and puncta(-) IDRs in organisms beyond humans.

## Results

### Establishing the condensation behavior of diverse IDRs

We previously compiled a database of 3,174 FO protein sequences and initially tested 166 of them for condensate formation in HeLa cells, with 96 of these forming condensates [puncta(+)] and 53 remaining diffuse [puncta(-)]. The sequences of the 96 puncta(+) FOs were significantly enriched in amino acids associated with protein disorder and in physicochemical features associated with phase separation, including the likelihood of pi-pi and pi-cation interactions and the presence of prion-like domains^34,36^. These observations led us to hypothesize that IDRs contribute to FO condensation behavior and that this set of 96 puncta(+) and 53 puncta(-) FOs (149 total) could serve as a source of both puncta(+) and puncta(-) IDRs.

We used the SAK pipeline^34^, which utilizes IUPred2A^37^, and Metapredict^38^, to identify 215 unique IDRs with long disordered regions (≥60 amino acids in length)^39^ within 83 puncta(+) and 47 puncta(-) FOs (see Methods). We note that 19 of the 149 puncta(+) and puncta(-) FOs did not display IDRs of sufficient length for inclusion. We expressed GFP-tagged forms of the 215 IDRs in HEK293T cells and scored them for condensate formation using established procedures^34^, including use of the PunctaTools image analysis pipeline^40^ (Supplementary Fig. 1a, b). IDRs that formed condensates in ≥24% of the imaged cells were scored as puncta(+); those that formed condensates in <24% of the imaged cells, or were diffuse in all cells, were scored as puncta(-) (Supplementary Fig. 1c, d). Some IDRs localized within nucleoli or other cellular structures lacking hallmark condensate features (e.g., round appearance with varied size) and were scored as nucleolar or other, respectively (Supplementary Fig. 1e, f), and not included in subsequent analyses. Of the 215 IDRs examined, 41 were classified as puncta(+), 137 as puncta(-), 15 as nucleolar, and 22 as other (Fig. 1a). Among the puncta(+) IDRs, 61% (25) formed condensates within nuclei, 10% (4) within the cytoplasm, and 29% (12) within both cellular compartments (Supplementary Fig. 1c, d) and the number and size of these condensates varied widely (Supplementary Fig.1c). Puncta(-) IDRs also exhibited varied sub-cellular localization (Supplementary Fig.1d) with many displaying diffuse fluorescence similar to that of GFP, the negative control (Supplementary Fig.1g).

**Fig 1.**
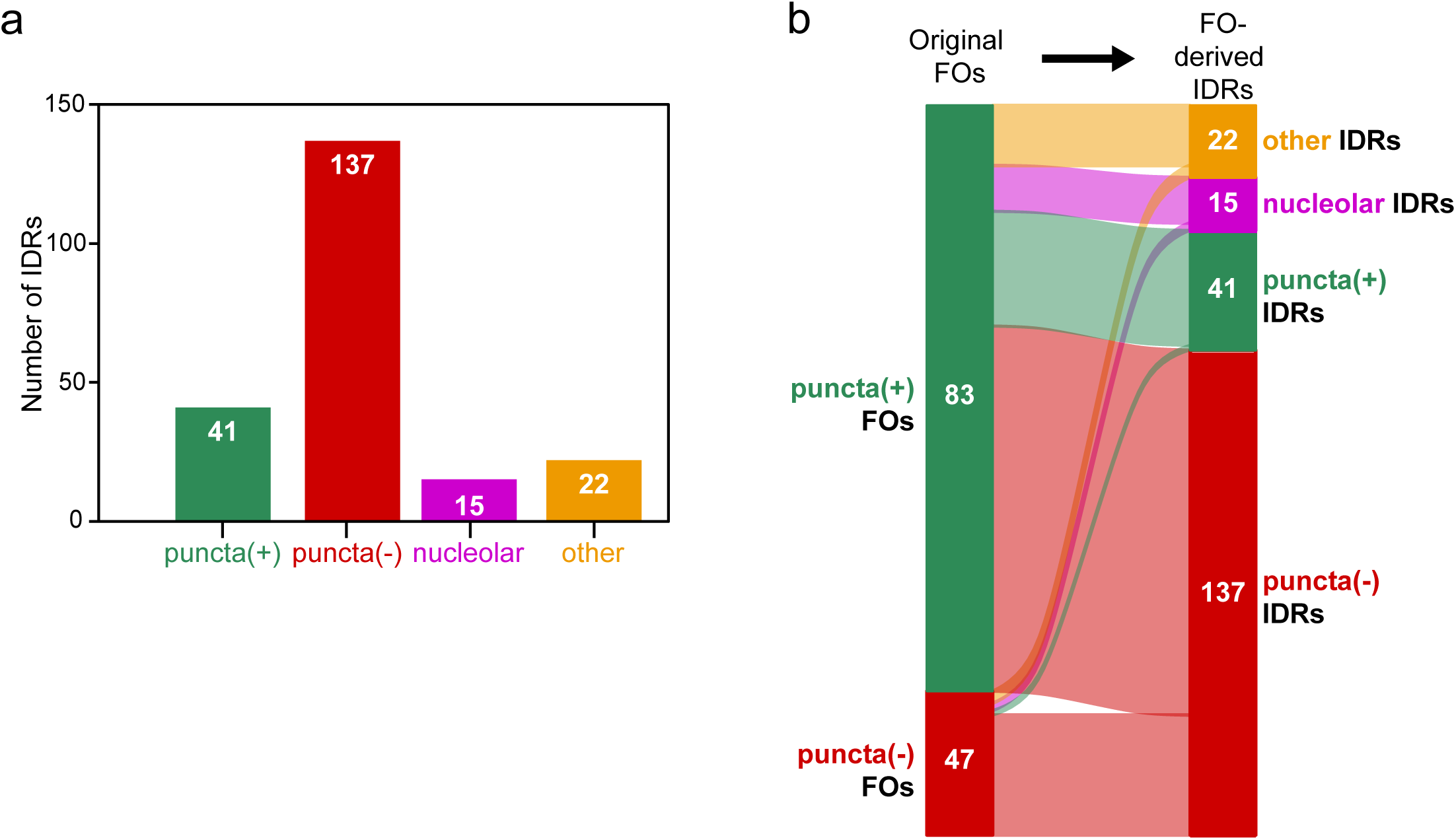
Condensate formation results from live cell imaging of mEGFP-tagged IDRs. **(a)** Quantification of the number of IDRs classified as puncta(+), puncta(-), nucleolar, or other (left). See “Methods” for details of these classifications. **(b)** Alluvial plot illustrating the puncta status of FOs and the IDRs derived from their sequences [puncta(+), green; puncta(-), red; nucleolar, magenta; and other, orange].

Our results showed that 48% of IDR-containing, puncta(+) FOs (40 of 83) display at least one puncta(+) IDR (Fig. 1b), leading us to hypothesize that puncta(+) IDRs contribute to condensate formation by these FOs. We tested this idea through IDR deletion analysis of five puncta(+) FOs, each containing a single puncta(+) and multiple puncta(-) IDRs (Supplementary Fig. 2a). We observed loss of condensate formation for four of the five FO IDR deletion mutants (Supplementary Fig. 2b), supporting that IDRs contribute substantially to condensate formation by these puncta(+) FOs. For the remaining IDR-containing, puncta(+) FOs (43 of 83) that lack a condensate-forming IDR, condensate formation is likely influenced by additional factors, such as synergy between IDRs and folded domains, or recruitment as clients into condensates formed by endogenous cellular biomolecules. Of the 47 puncta(-) FOs, one contained a puncta(+) IDR, and another contained an IDR that localized within nucleoli (Fig. 1b). For these two puncta(-) FOs, the condensate-associated IDRs they contain are insufficient to drive condensate formation by the full proteins, potentially due to solubilizing effects of folded domains and/or other IDRs in their sequences.

Overall, our results show that 19% (41 of 215) of the IDRs we tested form cellular condensates, with all but one of these derived from condensate-forming FOs. Most of the tested IDRs (64%, 137 of 215) scored as puncta(-), with many of these derived from condensate-forming FOs. These results indicate that autonomously puncta(+) IDRs contribute to condensate formation by about half of the FOs that exhibit this behavior, and that autonomously puncta(-) IDRs require synergy with other protein regions for the other half of condensate-forming FOs. Our previous studies of FOs indicated that sequence-derived physicochemical features are statistically accurate indicators of their condensation behavior. Based on these findings, we next asked whether these types of physical chemistry-based features could distinguish between puncta(+) and puncta(-) IDRs.

### Physicochemical features and amino acid enrichments governing IDR condensate formation

We analyzed the amino acid sequences of the 41 puncta(+) and 137 puncta(-) IDRs to identify physicochemical features associated with autonomous cellular condensate formation (Fig. 2a). For each sequence, we calculated the values of 600 diverse features, including PS-relevant physicochemical features (38 features from the SAK pipeline^34^), numerical indices related to amino acid physicochemical and biochemical properties (553 features from AAindex v9.2^41^ grouped into 12 classes; Supplementary Fig. 3), and molecular interaction-based features (9 features from LLPhyScore^21^). Feature values are reported as z-scores with respect to average values for the human IDRome (all IDRs ≥60 amino acids in length in the human proteome; see Methods). Amongst these 600 features, we identified 38 with average z-score values that were significantly different between puncta(+) and puncta(-) IDRs. To minimize redundancy, we removed 13 features that exhibited high mutual information (MI) scores with others (MI > 0.5) (Fig. 2a). The remaining 25 features sample a range of amino acid sequence-derived properties, including prion-like domain content, fraction of aromatic amino acids, potential for interactions, disorder content, charge content and patterning, and hydrophobic characteristics, which are differentially enriched or depleted in puncta(+) versus puncta(-) IDRs (Fig. 2b and Supplementary Dataset). Interestingly, secondary structure-related terms associated with extended conformations, e.g., sheets, and coils, are enriched in puncta(+) IDRs while those associated with compact conformations, e.g., helices and turns, are depleted. This collection of 25 features defines the physicochemical properties that, on average, underpin condensate formation by IDRs in the crowded, heterogeneous cellular environment.

**Fig 2.**
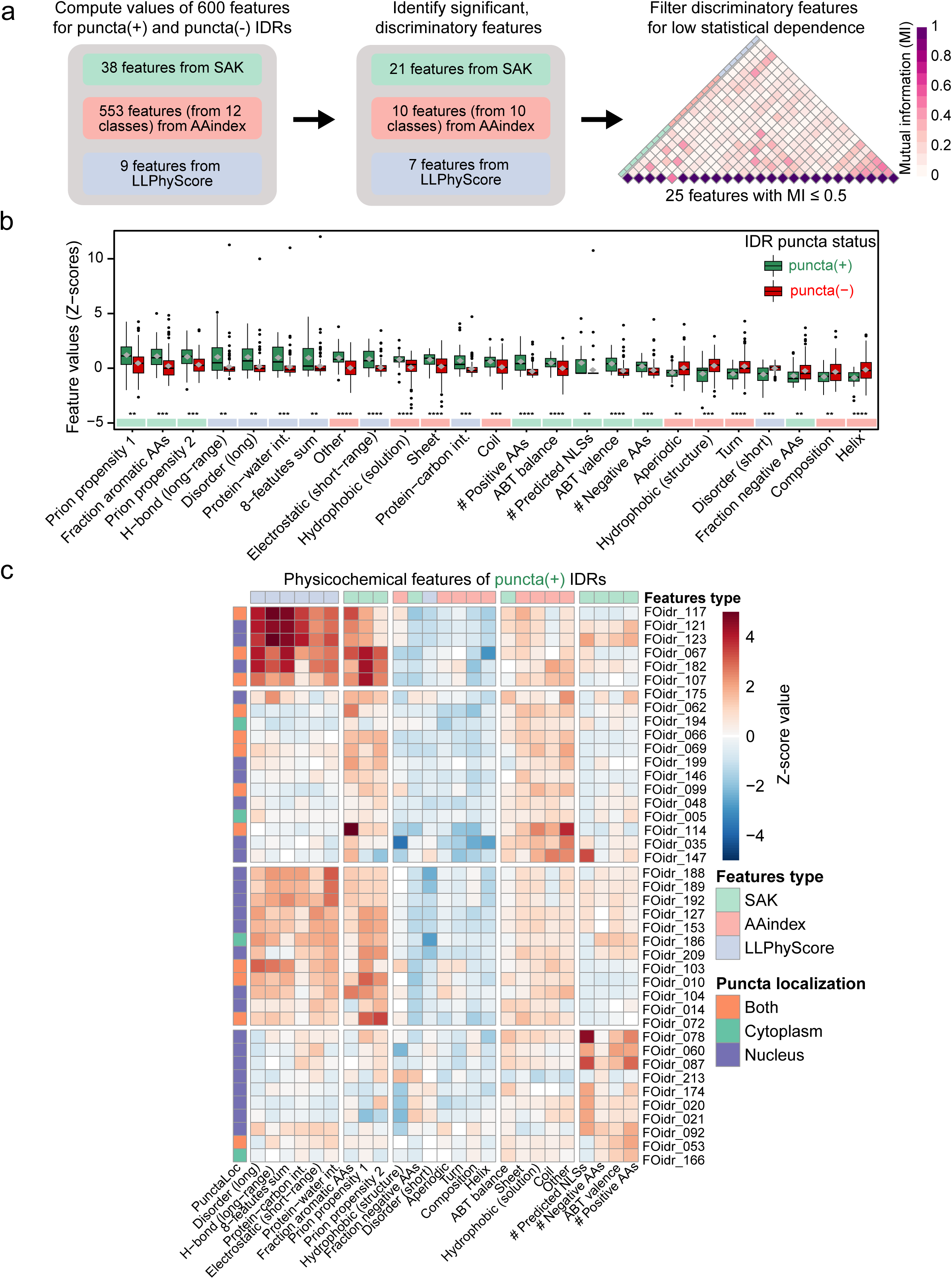
Physicochemical features differentiate between puncta(+) and puncta(-) IDRs. **(a)** Workflow of selection of the 25 most significant and non-redundant physicochemical features for 41 puncta(+) and 137 puncta(-) IDRs from SAK (cyan), AAindex database (red) and LLPhyScore (blue) (see Methods). To remove redundant features, the mutual information (MI) value of ≤ 0.5 was selected. See Supplementary Dataset for additional information on the physicochemical features used in these analyses. **(b)** Quantification of the enrichment or depletion of the 25 non-redundant, most significant physicochemical features for 41 puncta(+) and 137 puncta(-) IDRs with respect to the IDR sequences in the human proteome (human IDRome). Feature values for puncta(+) and puncta(-) IDRs are reported as z-scores using box plots in green (left) and red (right), respectively, along the *y*-axis. Each box shows the quartiles of the dataset, where the first black horizontal line of the box is the first quartile (25% of the data), the second black horizontal line is the second quartile or median (50% of the data), the third black horizontal line is third quartile (75% of the data). The whiskers extend to points that lie within 1.5 IQRs (interquartile range) of the lower and upper quartile and outliers are displayed as filled circles in black. The mean values of each feature are shown as diamond shapes in grey inside the boxes. Significance of the difference between the puncta(+) and puncta(-) IDRs was assessed using a two-sided Welch’s *t*-test and no adjustments were made for multiple comparisons (\**p*D<D0.05; \*\**p*D<D0.01; \*\*\**p*D<D0.001; \*\*\*\**p*D<D0.0001). The names of the physicochemical features are given at the bottom. The colored bars above the feature names represent feature types (SAK, light green; AAindex, light red; LLPhyScore, light blue). **(c)** Results of two-dimensional (2D) hierarchical clustering of the 41 puncta(+) IDRs based on the 25 most discriminatory physicochemical features as z-scores (columns) with respect to the human IDRome (see Methods) into four groups (Groups 1–4). The top row represents feature types (color scheme as in panel b). IDR feature values are color-coded in the rows, with IDR names are given on the right. The first column (left) represents the cellular localization of the IDR puncta (nucleus, blue; cytoplasm, green; or both, orange). The names of the physicochemical features are given at the bottom.

We next used two-dimensional (2D) hierarchical clustering to determine whether the individual IDRs within the puncta(+) and puncta(-) sets exhibited similar or different features and if they clustered into groups with similar features. The results showed that both sets were divided into four groups of IDRs, with members of the groups exhibiting similar patterns of physicochemical features (Fig. 2c, Supplementary Figs. 4, 5). For the puncta(+) IDRs, Groups 1 and 3 (with 6 and 12 IDRs, respectively) are most highly enriched in features reporting on different types of molecular interactions (6 features from LLPhyScore), and on aromatic residue content and prion-like domain content (3 features from SAK). Puncta(+) IDR Group 2 (with 13 IDRs) is weakly enriched in features related to aromatic residue content and prion-like domain content, and others reporting on charge balance (from SAK) and β-sheet and coil secondary structure content (from AAIndex). In contrast, puncta(+) IDR Group 4 (with 10 IDRs) exhibits moderate enrichment of charge related features (4 features from SAK) and otherwise mixed and weak feature enrichments and depletions.

Average, group-wise feature enrichments for puncta(+) IDRs were accompanied by distinct patterns of average amino acid enrichments (see Methods), variably including enrichment of phenylalanine (Group 1), tyrosine (Groups 1-3), glycine, asparagine and glutamine (Group 1), and arginine and lysine (Group 4; Supplementary Fig. 5a, b), which are known from prior studies to be enriched in condensate-forming proteins^13,15,17,34,42^. The puncta(-) IDR groups exhibited feature and amino acid enrichments that were either weaker than (Groups 1’ and 4’) or different from (Group 3’) those of the puncta(+) IDRs with the exception of puncta(-) Group 2’, whose enrichments resemble those of puncta(+) IDR Group 3. However, the magnitude of the feature enrichments and depletions for Group 3 puncta(+) IDRs are generally greater than those for Group 2’ puncta(-) IDRs (encoded as the red and blue color intensities in the feature heatmaps; Fig. 2c, Supplementary Figs. 4), suggesting that subtle differences in physicochemical feature profiles govern the condensation behavior of IDRs in these two groups. In summary, our results establish the physicochemical feature and amino acid enrichment landscape of IDRs that do and do not form condensates in HEK293T cells.

### Predicting IDR cellular condensation using physicochemical features and machine learning

We next asked whether our IDR dataset could be leveraged to accurately predict the condensation behavior of additional, experimentally untested IDRs. Using 25 physicochemical features (Fig. 2b) for 41 puncta(+) and 137 puncta(-) IDRs as training data (termed the Training IDRs), we applied H_2_O AutoML^43^ to evaluate 120 machine learning (ML) models for prediction of cellular IDR condensation behavior (Fig. 3a). A stacked ensemble model comprised of three tree-based models was superior among the tested models (termed the IDR-Puncta ML model, see Methods) and displayed the following performance metrics based on 25-fold cross validation: AUC [area under the ROC (receiver operating characteristic) curve], 0.98; AUCPR (area under the precision-recall curve), 0.93; accuracy, 0.95; and balanced accuracy, 0.92 (Fig 3b). Analysis using Shapley Additive exPlanations (SHAP)^44^ showed that diverse physicochemical features contribute to predictions of IDR condensate formation (see Methods), including those reporting on charge-, disorder-, and hydrophobicity-related properties (Supplementary Fig. 6a). We independently verified IDR-Puncta ML model performance using 30 Human IDRs out of the total 33 IDRs (Verification IDRs) selected to have low degree of sequence identity based on pairwise alignment (<55% for one sequence and <20% for the remaining 32 sequences) with the Training IDRs (Fig. 3c; see Methods), and observed accuracy similar to that obtained during cross validation [AUC, 0.95; AUCPR, 0.88; accuracy, 0.90; and balanced accuracy, 0.92 (Fig. 3d; Supplementary Dataset)]. We excluded three verification IDRs from the ML model performance evaluation, which were experimentally categorized as either “nucleolar” or “other”. We next performed dimensionality reduction analysis using the 25 physicochemical features of the Training IDRs and the IDRs (≥60 amino acids in length) from human proteome combined. The analysis revealed that the physicochemical features of the FO-derived Training IDRs we tested spanned the feature landscape of the human IDRome (Supplementary Fig. 6b). This result further confirms that the IDR-Puncta ML model can be applied to the human IDRome to understand the prevalence of puncta(+) IDRs in human proteins. The human proteome (20,396 proteins) contains 12,899 IDRs (derived from 8,067 proteins, see Methods) and 1,572 of these (12%) were predicted to be puncta(+) using the IDR-Puncta ML model (Fig. 3e, Supplementary Fig. 6c), indicating that the propensity for condensate formation is a specialized property of human IDRs.

**Fig 3.**
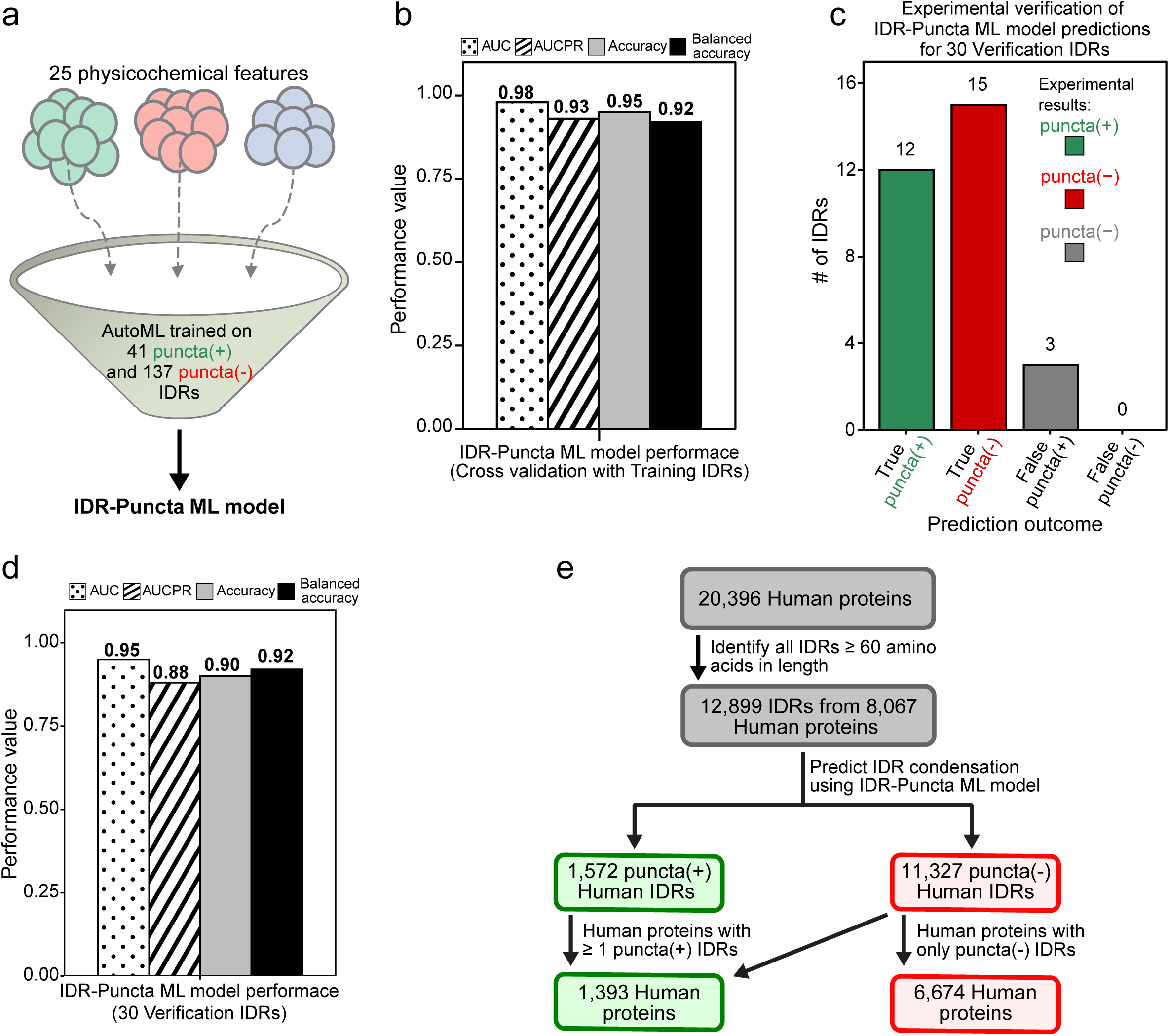
A Machine Learning model for predicting condensate formation probability of IDRs. **(a)** Supervised Machine Learning was used to develop a Stacked Ensemble model (termed IDR-Puncta ML model; see Methods) trained using the 25 low mutual information physicochemical features from SAK (9 features; cyan), AAindex (9 features; red) and LLPhyScore (7 features; blue) for 41 puncta(+) and 137 puncta(-) IDRs (termed Training IDRs). **(b)** Performance metrics [area under the curve (AUC, dots), area under the precision-recall curve (AUCPR, stripes), accuracy (grey), and balanced accuracy (black)] for the IDR-Puncta ML model using 25-fold cross validation (CV) with the Training IDRs. **(c)** Results of prediction of condensate formation behavior [x-axis, puncta(+) or puncta(-)] using the IDR-Puncta ML model for the 30 puncta(+) and puncta(-) Verification IDRs out of the total 33 Verification IDRs (see Methods). The number of predicted true puncta(+) and true puncta(-) IDRs from the IDR-Puncta ML model are shown as green and red bars, respectively. The number of predicted false puncta(+) IDRs is shown as a grey bar. No false puncta(-) IDRs were predicted. **(d)** Performance metrics [area under the curve (AUC, dots), area under the precision-recall curve (AUCPR, stripes), accuracy (grey), and balanced accuracy (black)] for the IDR-Puncta ML model using the 30 Verification IDRs. **(e)** Predicted condensation behavior of all IDRs in human proteome, termed the human IDRome (12,899 IDRs, in total), using the IDR-Puncta ML model.

### Human proteins with puncta(+) IDRs are enriched for RNA processing-related functions

Sequence mapping of the ML prediction results showed that 1,393 human proteins contain one or more puncta(+) IDRs [some of these proteins also contain one or more puncta(-) IDRs] while 6,674 proteins contain only puncta(-) IDRs (Fig. 3e). We next asked whether the human proteins that contain one or more puncta(+) IDR(s) are associated with specific biological functions using Gene Ontology (GO) biological process enrichment analysis in comparison with all human proteins. A similar analysis was performed for human proteins containing only puncta(-) IDRs. Three of the top four most highly and significantly enriched parent biological process terms for proteins with puncta(+) and puncta(-) IDRs were similar (e.g., positive regulation of transcription, cell division, and actin cytoskeleton regulation; Fig. 4a), indicating that functions related to these processes are common to proteins with IDRs regardless of condensate formation. In contrast, the most highly enriched parent term for puncta(+) IDRs was RNA processing (3.5-fold enrichment, Fig. 4a), which aggregates numerous enriched child terms related to regulation and processing of RNA, including metabolism and splicing (Supplementary Fig. 7a). We note that the magnitude of functional term enrichments was greater for proteins with puncta(+) than puncta(-) IDRs (Fig. 4a, Supplementary Fig. 7a). These results indicate that proteins involved in RNA-related processes are enriched in IDRs prone to condensate formation and suggest that these processes occur within condensate environments. Analysis of GO cellular component terms, which report on sub-cellular localization, showed that proteins with puncta(+) IDRs were enriched for terms including nucleolus, nuclear body, nuclear speck, nuclear protein-containing complex, and spliceosomal complex (Fig. 4b, Supplementary Fig. 7b), supporting the association of these proteins with several types of nuclear biomolecular condensates. The condensates associated terms (e.g., nucleolus, nuclear body, and nuclear speck) were not enriched in human proteins with puncta(-) IDRs (Fig. 4b, Supplementary Fig. 7b).

**Fig 4.**
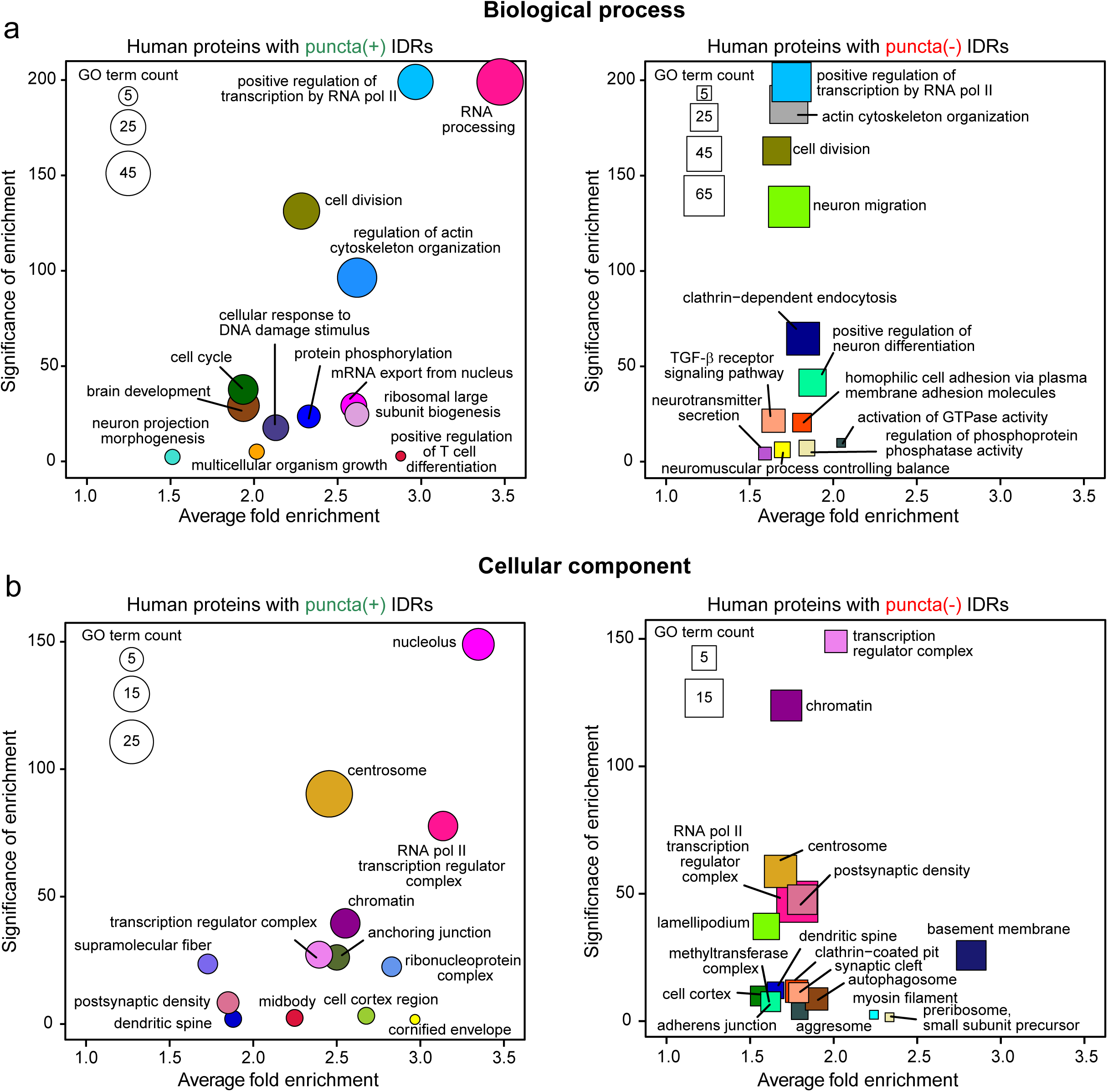
Biological functions of human proteins with puncta(+) and puncta(-) IDRs. Scatter plots showing Gene Ontology (GO) enrichment analysis in two categories, biological processes (a) and cellular component (b). The y-axis shows the combined adjusted *p*-value (p.adj) of the enriched GO terms, and the x-axis gives the average fold-enrichment after grouping the GO terms using semantic similarity analysis (see Methods). **(a)** Enriched biological processes for proteins with predicted puncta(+) IDRs (left) and proteins with only predicted puncta(-) IDRs (right). **(b)** Enriched cellular components for proteins with predicted puncta(+) IDRs (left) and proteins with only predicted puncta(-) IDRs (right). Symbols (a, b) are color coded according to the different grouped GO terms from semantic similarity analysis and the symbol size is proportional to the number of GO terms in each group. For the grouped GO terms combined significance was obtained using Fisher’s method and average fold enrichment was computed using log transformation. The human proteome was used as background for all the GO term analysis (see Methods).

In summary, our IDR-focused analyses indicate that 40% of human proteins (8,067 of 20,396 proteins) exhibit one or more IDRs ≥60 amino acids in length and that these are enriched for functions associated with transcription, cell division, and actin cytoskeleton. The IDR-containing human proteins with potential for condensate formation [e.g., displaying puncta(+) IDRs; 1,393 of 8,067 proteins] are uniquely enriched in functional terms involving RNA-related processes and in localization terms associated with nuclear membraneless organelles. While others have associated condensate formation by proteins with RNA-related biological processes^45–47^, our findings extend these results by showing through large-scale, unbiased analyses that human proteins with condensation-prone IDRs are highly enriched for RNA-related biological functions and localization within nuclear membraneless organelles.

### Proteins with puncta(+) IDRs are enriched in membrane-less organelles (MLOs)

We independently tested our findings on preferential localization of human, puncta(+) IDR-containing proteins within biomolecular condensates by examining their occurrence in the membraneless organelle (MLO) protein database, PhaSepDB^48^, which compiles information on proteins demonstrated to undergo PS and/or reported to be associated with one or more MLOs. We retrieved 499 human MLO-associated proteins from PhaSepDB and identified 345 with one or more IDRs ≥60 amino acids in length. We also compiled a control dataset comprised of membrane associated proteins, which we reasoned do not autonomously form biomolecular condensates. Specifically, we examined proteins found at membrane-bound organelle contact sites (MCSs; compiled in MCSdb^49^), wherein membranes of two different organelles are in close proximity but do not fuse^50^. We obtained 199 human proteins from MCSdb and identified 90 with one or more IDRs. To enhance the robustness of our analyses, we only included proteins annotated with high confidence; our final MLO and MCS datasets included 345 and 90 human proteins, respectively, with only 2 proteins common between the two datasets (Supplementary Fig. 8a, see Methods). We next applied our IDR-Puncta ML model to the two protein sets and determined that 134 of 345 MLO proteins (39%) and 5 of 90 MSC proteins (6%), respectively, contain one or more IDRs predicted to form condensates (Fig. 5a). In comparison, we reported above that 1,393 of 8,067 human IDR-containing proteins (17%) contain puncta(+) IDRs (Fig. 3e). These results, showing that proteins with puncta(+) IDRs are much more highly enriched in MLO proteins than in MCS proteins or human IDR-containing proteins support our independent findings based on analysis GO cellular component localization terms (Fig. 4b, Supplementary Fig. 7b). While the occurrence of a puncta(+) IDR within the sequence does not necessarily indicate that a protein will autonomously form condensates, it is highly suggestive of localization within an MLO. The 134 MLO proteins with puncta(+) IDRs are predominantly associated with nuclear MLOs although some are associated with cytoplasmic MLOs, including stress granules and P-bodies (Fig. 5b). These proteins are most frequently associated with nuclear speckles, nucleoli, and paraspeckles, MLOs involved in different RNA-related processes, consistent with our observation that the functional term, RNA processing, was the most highly enriched amongst human proteins with puncta(+) IDRs (Fig. 4a, Supplementary Fig. 7a). These findings highlight a strong correlation between proteins containing puncta(+) IDRs and their preferential localization within MLOs, particularly those with roles in RNA-related processes. This relationship underscores the importance of condensate-driven compartmentalization in organizing and regulating RNA metabolism, indicating how MLOs support complex biochemical processes.

**Fig 5.**
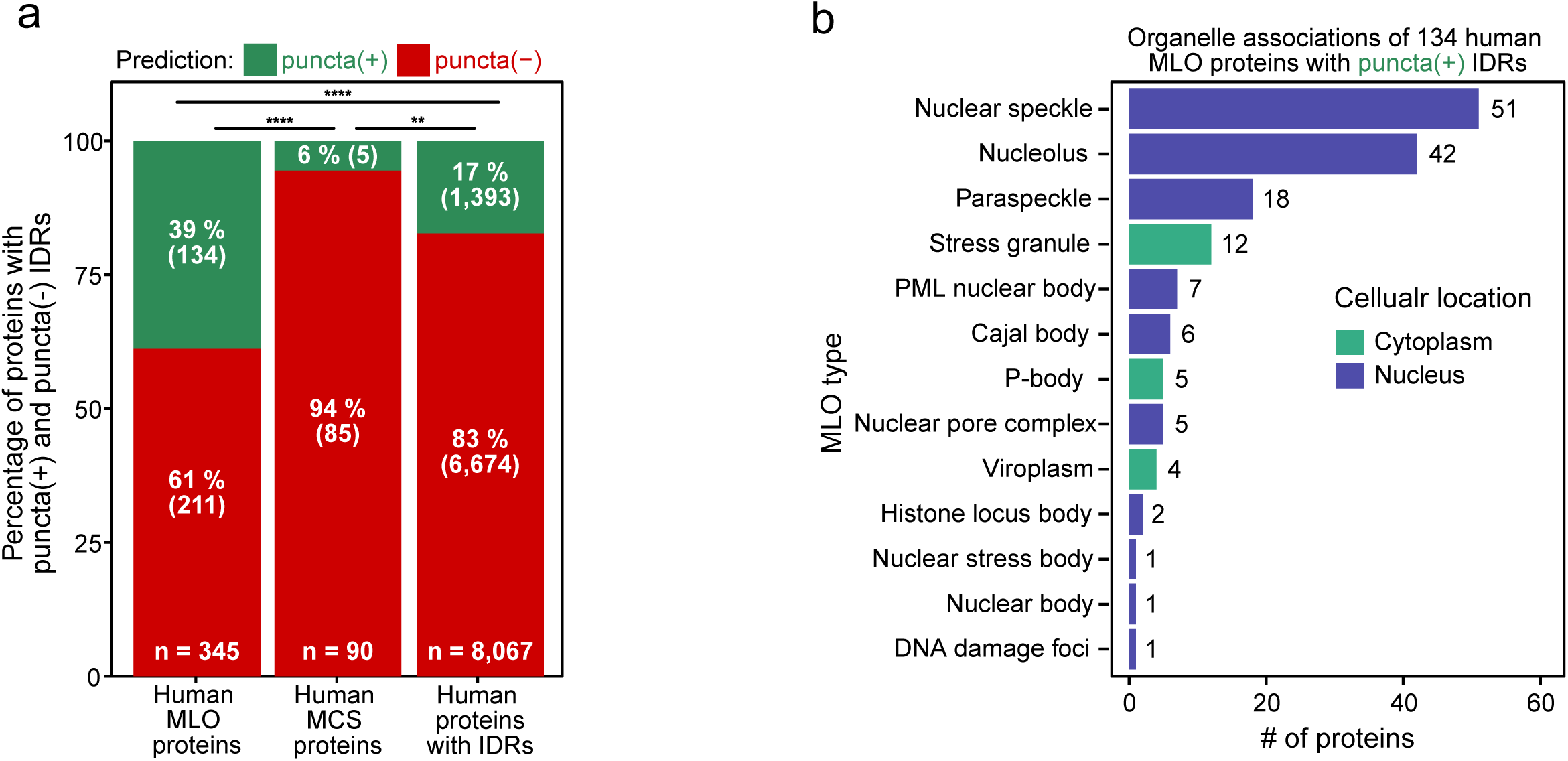
Prevalence of condensate prone IDRs in MLOs. **(a)** Quantification of the presence of puncta(+) or only puncta(-) IDRs in MLO proteins (left bars), MSC proteins (middle bars) and human proteins (right bars). The puncta(+) or only puncta(-) status of IDRs in the sets of proteins was determined using the IDR-Puncta ML model [green, percentage of proteins with at least one punta(+) IDR; red, percentage of proteins with only punta(-) IDRs]. Significance was assessed using Fisher’s exact count test and no adjustments were made for multiple comparisons (\**p* < 0.05; \*\**p* < 0.01; \*\*\**p* < 0.001; \*\*\*\**p* < 0.0001). **(b)** MLO localization of proteins from the MLO set containing at least one predicted puncta(+) IDR. The colors of the bars represent the two major cellular compartments (nucleus, blue; cytoplasm, green) and the numerical values above the bars indicate number of proteins along the x-axis.

## Discussion

Protein PS plays a key role in cellular organization and function^6,13^. The ability to accurately predict PS propensities of proteins is critical to understand cellular compartmentalization and its roles in biology and disease. Recent advances fostered the development of several computational tools, each leveraging different approaches to analyze protein PS^21,23,30,31^. However, the lack of standardized PS testing methods and robust PS negative datasets introduces implicit and difficult-to-measure biases among these different prediction methods^33^. Furthermore, many of these predictors use literature-mined datasets from different databases, which often exhibit inconsistencies and uncertainty due to non-standardized experimental conditions. We previously developed the FO-Puncta ML model to predict the cellular condensation behavior of FOs^34^. However, FOs differ from human proteins due to their unique sequence features, which result from aberrant gene translocation. As a result, the model is not optimized to recognize the broader sequence-based features associated with phase separation-prone human proteins. Additionally, this model is designed to assess the condensation behavior of full-length proteins and does not identify which specific region(s) of the FO drives this behavior. Since many proteins harbor multiple IDRs, only some of which may contribute to PS, identifying the exact IDR(s) responsible for condensate formation is crucial. Such specificity not only illuminates the molecular basis of PS but also enables targeted sequence modifications. An improved understanding of key features of PS prone IDRs such as amino acid composition, charge distribution, and potential for multivalent interactions is essential for rationally controlling the phase behavior of proteins.

Building on these advances, we developed the IDR-Puncta ML model, a machine-learning tool that accurately predicts the condensate-forming potential of IDRs, based on sequence-derived physicochemical features. The IDR-Puncta ML model was trained on experimentally tested IDRs using standardized cellular assays and does not report on the specific type (homotypic or heterotypic) of interactions that may drive condensate formation. Our model addresses limitations that may have variably impacted existing tools, such as unvalidated datasets, lack of cellular context, potential overfitting due to feature redundancy, and narrowly focused features. We experimentally validated the IDR-Puncta ML model using randomly selected IDRs from the human proteome, confirming its accuracy and broad applicability.

Recent work by Bülow, et al., predicts that 5% of IDRs from the human proteome can undergo homotypic PS under physiologically relevant conditions (e.g., sequences with transfer free energy values (ΔG^tr^) less than −2k_B_T^35^). We applied this model to our set of 41 puncta(+) IDRs and found that Group 1 IDRs are predicted to undergo homotypic phase separation, with a mean ΔG^tr^ value of −4.89 k_B_T (Supplementary Fig.9a). Group 3 IDRs have a mean ΔG^tr^ value of −1.87 k_B_T, indicating a moderate tendency toward homotypic phase separation. Conversely, IDRs from Group 2 (mean ΔG^tr^ = −0.82 k_B_T) and Group 4 (mean ΔG^tr^ = −0.29 k_B_T) are less likely to undergo homotypic PS (Supplementary Fig.9a). These trends align with the distinct sequence feature enrichment observed in each group (Supplementary Fig. 4a). Groups 1 and 3 show similar feature enrichment patterns, although the enrichment is less pronounced in Group 3. Groups 2 and 4 display unique feature enrichment profiles, distinguishing them from Groups 1 and 3. These results highlight variability in PS propensities across different IDR groups. We hypothesize that puncta(+) IDRs, exhibiting distinct patterns of physicochemical features and amino acid enrichments (e.g., IDRs in Groups 1-4; Fig. 3a), will display different conformational properties and engage in various intra- and inter-polypeptide chain interactions, both homotypic and heterotypic, that drive condensate formation. However, testing this hypothesis requires further investigation in the future.

We explored the biological implications of protein condensate formation by conducting a detailed functional annotation of proteins containing condensate-forming IDRs. Using our predictive model, we identified 1,393 proteins containing at least one predicted puncta(+) IDR (∼17% of all IDR-containing human proteins). These findings align with studies suggesting that PS driven by IDRs alone is limited, with additional factors such as folded domains influencing condensate formation^25,51^. Our data indicates that proteins with predicted puncta(+) IDRs are highly enriched in RNA-related biological processes, specifically RNA-processing and splicing. It is increasingly evident that RNA-related processes (including transcription, splicing, and translation) are frequently associated with condensate formation^52–54^. Furthermore, the proteins with puncta(+) IDRs are enriched in nuclear speckles, nucleoli, and paraspeckles, MLOs known to facilitate various RNA-related processes. These data serve as a valuable resource for hypothesis-driven research into the roles and mechanisms of condensate formation by IDRs in RNA biology, by providing insight into patterns of physicochemical features in IDRs and their cellular condensation behavior.

Our study establishes a framework for examining the link between IDR mediated protein condensation and biological function. Herein, we present a valuable resource for mapping the potential of human IDRs to form condensates in cells, offering insights into their physicochemical features, and associated biological functions, especially in the context of PS. Our prediction model should be useful in guiding cellular experiments to explore the role of novel IDRs in PS. Our model may also be used in providing key insights into condensate pathology. For example, aberrant condensate formation has been associated with several human diseases, especially in neurodegeneration and cancer^4,7^. Identifying specific IDR that drives pathological condensate formation can aid in finding the association between PS and disease progression. As our model was trained on FO-derived IDRs, it is particularly suitable to analyze the cancer proteome. Identifying puncta-forming IDRs in cancer-associated proteins will provide critical insights into their possible oncogenic mechanism. The IDR-Puncta ML holds significant potential in interpreting the impact of disease-associated mutations in IDRs that alter cellular PS behavior for understanding complex diseases, including neurodegenerative disease and cancer^5,55^. Beyond individual IDR, our model can also be applied to a proteome-wide assessment of IDR-mediated PS across different organisms to explore the prospect of PS being an evolutionary conserved mechanism in regulating diverse cellular processes and stress response^56^. The IDR-Puncta ML model can also be applied in synthetic biology and biomaterials engineering. Our work provides refined knowledge of sequence features associated with condensate-prone IDRs and tools to design and modify PS-prone IDRs, which will assist the engineering of synthetic protein condensates and new biomaterials with tunable phase-separating properties^57,58^.

### Limitations of our study

We note a few factors to be considered when evaluating our findings and conclusions. First, the training set for our model used IDRs from a subset of FOs, which may limit the generalizability of our findings to other proteins or IDRs with different characteristics. However, our analyses showed that the physicochemical features of the FO-derived IDRs we tested spanned the feature landscape of the human IDRome, mitigating this concern. Second, we focused on IDR-driven condensate formation of proteins and excluded PS promoted by folded domains (oligomerization domains, nucleic acid binding domains)^59,60^ as well as PS facilitated by multiple IDRs working in concert. These factors limit our ability to identify all condensation-prone proteins in the human proteome. Moreover, we segregated IDRs that flank a folded domain and tested them individually, potentially overlooking the enhanced effects of multiple IDRs acting together. Additionally, while we identified physicochemical feature patterns linked to puncta(+) behavior in IDRs, we did not explore how these patterns affect IDR conformational dynamics or multivalent interactions, both crucial for condensate formation.

## Methods

### IDR selection for in-cell expression

IDRs were identified by analyzing the 149 Fusion oncoprotein sequences in our previous work^34^ using sak.stjude.org, which identified IDRs based on continuous stretches of disordered residues ≥ 60 residues according to the IUPRED2A algorithm^37^. Sequences were also analyzed with Metapredict (version: V2)^38^, and Metapredict identified IDRs were added to the IDR database when there was not an analogous IUPRED2A identified IDR. In some cases, IUPred2A and Metapredict identified multiple adjacent IDRs that did not form puncta. In these cases, combined IDRs that contained all the adjacent disordered regions were created and tested. A total of 215 unique IDRs were identified among which 188 were predicted using SAK, 23 with Metapredict, and 4 IDRs were identified by both IUPred2A and Metapredict.

### Cloning

We previously reported *Escherichia coli* codon optimized full-length plasmids for cellular expression of FOs^34^. IDRs and FO-IDR deletion mutants were generated using PCR with the respective FOs as a templates. PCR reactions were performed using Q5 High Fidelity 2x Mastermix and primers were designed with 5’-Not1 and 3’-Xba1 restriction sites. PCR fragments and the destination vector (CL20) were cut with Not1 and Xba1 restriction enzymes and ligated using New England BioLabs’ Quick Ligation Kit per the manufacturer’s instructions. Ligation reactions were transformed into chemically competent bacteria (NEB 5-alpha). All plasmid sequences were confirmed by whole plasmid sequencing.

### Cell culture and transient transfections

HEK293T cells (ATCC; RRID: CVCL_0063) were cultured in DMEM with high glucose (Gibco) and supplemented with 1× penicillin/streptomycin (Gibco), 10% FBS (HyClone), and 6 mmol/L l-glutamine (Gibco) and maintained at 37 °C in 5% CO_2_. Cells were tested for Mycoplasma every 2 months using PCR (e-Myco plus, LiLiF). Cells were authenticated by short tandem repeat profiling (PowerPlex Fusion at the St. Jude Hartwell Center). Cells were transfected in a 96-well plate with 100 ng of plasmid DNA in the CL20 vector backbone using FuGENE HD (Promega) per the manufacturer’s instructions. All IDRs were N-terminally tagged with monomeric EGFP (A207K mutation in EGFP), and EGFP was used for the empty vector control plasmids. Cells were used for a maximum of 25 passages after thawing.

### Confocal microscopy imaging

All microscopy images were acquired on a 3i Marianas system (Denver, CO) configured with a Yokogawa CSU-W spinning disk confocal microscope utilizing a 100x Zeiss objective, 405 nm (Hoechst) and 488 nm (mEGFP) laser lines, and Slidebook (RRID: SCR_014300) 6.0 (3i). 3D images of cells were captured as z stacks with 0.3 µm spacing between planes, spanning 12 µm in total. Live HEK293T cells were imaged at 37 °C in phenol red-free DMEM with high glucose (Gibco) supplemented with 1× penicillin/streptomycin, 10% FBS, 6 mmol/L l-glutamine, and 25 mmol/L HEPES.

### Quantitative image analysis to classify IDRs as puncta (+), puncta (-), Other, and Nucleolar

Fluorescent microscopy images were segmented and quantified using the PunctaTools pipeline^40^. In brief, cells were segmented using eGFP as the primary signal, and Hoechst as a secondary signal to establish nuclei. Cells were segmented as sets of 10 layers to minimize cell-segmentation errors, then combined into 3D stacks using the Cellpose algorithm^61^. Condensates (termed puncta here) were then segmented by filtering respective channels with a scale adapted Laplacian of Gaussian (LoG) filter, thresholding the result, and applying watershed segmentation using the maxima of the LoG filtered image as seeds. Using this method, cells were recorded as having puncta, or not. A threshold of 24% of expressing cells was set based on agreement with the manual assessment of puncta status. After quantification, IDR puncta status was manually verified by two independent researchers, where segmentation errors and alternate classifications (Other, Nucleolar) were assigned, after which final classifications were assigned to each IDR.

### Calculation of amino acid sequence-derived physicochemical features of IDRs

To understand the potential of condensate formation by IDRs, we deployed physicochemical features that were computed for each IDR amino acid sequence. Specifically, we obtained features from our previous work^34^ through the SAK pipeline (sak.stjude.org), the Amino Acid Index (AAindex) database^41^, and the recently developed LLPhyScore resource^21^. We accessed amino acid properties from the AAindex database, which is a curated set of numerical indices representing various physiochemical and biochemical properties of amino acids. Additional features were obtained from the LLPhyScore resource, which includes a predictor of IDR-driven phase separating proteins based on underlying physicochemical interactions, including, solvent contacts, disorder, hydrogen bonds, pi-pi and cation-pi contacts, electrostatic interactions, and secondary structure. The Python package *protlearn* (version: 0.0.3) was used to extract 553 amino acid properties from the AAindex database with no missing values that belong to 12 property classes^25^ (Supplementary Fig. 3). The features from LLPhyScore were computed using the standalone package from GitHub (github.com/julie-forman-kay-lab/LLPhyScore) based on the model trained on both folded proteins in the PDB and proteins from the human proteome.

### Human IDRome sequences

To represent the human IDRome, we first obtained all human Swiss-Prot (reviewed) proteins from Uniprot release 2023_04. Sequences were excluded from the collection if their length was less than 10 residues, or if any characters outside of the natural amino acids were present. This resulted in a total of 20,396 human protein sequences (termed the human proteome here). From the human proteome, we identified 13,047 IDR sequences (from 8,067 proteins) through sak.stjude.org with lengths ≥ 60 residues, which were termed the human IDRome.

### Calculation of amino acid enrichment for IDR sequences

Amino acid enrichment for each IDR sequence was calculated using equation 1,

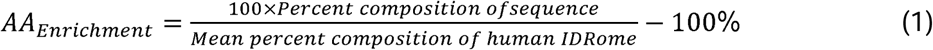

where “Percent composition of sequence” is the percent composition of a particular amino acid in the sequence being evaluated and “Mean percent composition of human IDRome” is the mean percent composition of a particular amino acid in a database of human IDR sequences (human IDRome). Amino acid compositions of the IDR sequences were computed using the *protr*^62^ package (version: 1.7.0) in R.

### Analysis of sequence-derived physicochemical features

First, we identified that for the 41 puncta(+) and 137 puncta(-) Expressed IDRs (collectively termed the Training IDRs; 178 IDRs, in total) values of the 38 features obtained from sak.stjude.org, values of the features, “ABT valence”, “ABT balance”, and “ABT density”, were missing for ∼5% of IDRs, whereas “PAPAprop” values were missing in ∼8% of IDRs, due to the limitation of the IDR length requirement for the calculation of these features. Therefore, we replaced all the missing values using the non-missing median values of these features from the puncta(+) and puncta(-) Expressed IDRs, respectively. We next performed the two-sided t-test for the 38 features from sak.stjude.org and identified 21 features that showed significant differences (p-value ≤ 0.01) between the 41 puncta(+) and 137 puncta(-) IDRs using the *rstatix* package (version: 0.7.2) in R (version: 4.2.1). Similarly, we identified 10 features from the AAindex database (with the most significant feature difference from each of the 12 classes) and 7 features from LLPhyScore (out of the total of 9 LLPhyScore features) that showed significant differences (p-value ≤ 0.01) between the 41 puncta(+) and 137 puncta(-) IDRs. Effect size was calculated using the *effsize* package (version: 0.8.1) (https://zenodo.org/record/196082) in R. Next, to identify interdependence of the 38 sequence-based physicochemical features for the 41 puncta(+) and 137 puncta(-) IDRs, we computed mutual information (MI) among these features using *infotheo* (version: 1.2.0.1) package in R. Features that displayed strong mutual dependence with others were removed, which resulted in 25 features with low MI (≤ 0.5). These 25 physicochemical features were converted to z-scores using the *scale* function in the R package, with respect to the human IDRome sequences. We next performed hierarchical clustering based on Euclidean distance and using the complete linkage method, as implemented in the *pheatmap* package (version: 1.0.12) in R, using the z-scores for the noted 25 features to identify groups within the 41 puncta(+) IDR set with related physicochemical features. For the 137 puncta(-) IDR set, we performed hierarchical clustering based on Manhattan distance and Ward’s minimum variance method, using the z-scores of the 25 physicochemical features.

### Supervised machine learning for puncta classification

We employed the automatic machine learning (AutoML) tool within the *h2o.ai* ^43^ (version: 3.44.0.1) package in R to classify the puncta (+) and puncta (-) IDRs and predict the probability of condensate formation using data for the 178 Training IDRs [consisting 41 puncta(+) and 137 puncta(-) IDRs from the Expressed IDRs]. Using the 25 sequence-based physicochemical features with low MI for the 178 Training IDRs, we set *nfolds*=25 for 25-fold cross-validation (CV) and generated 120 models from AutoML. Additionally, we set *include_algos* to the H2O tree-based models [Gradient Boosting Machine (GBM), Distributed Random Forest (DRF) including Extremely Randomized Trees (XRT), and Extreme Gradient Boosting (XGBoost)] and Stacked Ensembles, using otherwise default parameters in H2O AutoML. A “Best of Family” Stacked Ensemble Model consisting of three base models GBM, XRT, and DRF performed the best amongst the 120 models tested (termed the IDR-Puncta ML model). The Stacked Ensemble metalearner using an Elastic net regularized (with CV) Generalized Linear Model (GLM) resulted in the ML model with GLM coefficients, 0.62, 0.46, and 0.18 for the three base models GBM, XRT, and DRF, respectively. Performance of the model was based on the metrics logistic loss or cross-entropy loss (the difference between predicted probabilities and actual values) value of 0.17 for the 25-fold CV set with 178 Training IDRs. We determined a threshold value for the condensate formation probability of 0.40 based on a maximum accuracy value of 0.95, maximum F1 score (harmonic mean of the precision and recall) value of 0.89, and maximum absolute Matthew’s correlation coefficient value of 0.86 for classifying the puncta(+) and puncta(-) IDRs for the 25-fold CV set with 178 Training IDRs. After its establishment, we applied the IDR-Puncta ML model to 30 Verification IDRs [with experimentally determined puncta(+) and puncta(-) status] out of the total 33 Verification IDRs from human IDRome. We excluded three Verification IDRs which were experimentally determined as either “nucleolar” or “other” from the ML model performance evaluation. To measure the degree of similarity between the IDR sequences in the Training and Verification sets, we performed pairwise sequence alignment using protein BLAST (version: BLAST+ 2.14.1). We defined pairwise normalized alignment score, fraction of identical matches as,

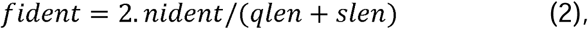

where *qlen* is the length of the query sequence, *slen* is the length of the subject sequence and *nident* is the fraction of identical matches between the query and subject sequences. *fident* varies betwee 0 and 1, with 0 indicating no identical residues between the query and subject sequence, and 1 indicating an exact match between the query and subject sequences. We used the packages *PRROC* (version: 1.3.1)^63^ in R to compute the performance metrics AUC and AUCPR for the Verification IDRs. We next applied Shapley Additive exPlanations (SHAP) analysis^44^ to determine the importance of the 25 physicochemical features to IDR-Puncta ML model predictions. SHAP is a game-theoretic approach to explain the output of any machine learning model. First, we computed Shapley values (*SHAP_value_*) of the features for each of the 178 Training IDRs using the *h2o.predcit_contribution* function in the *h2o.ai* package in *R.* Next, we computed SHAP importance for each feature from the base GBM model with highest contribution to the IDR-Puncta ML model as,

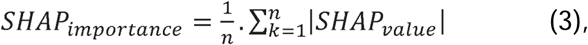

where *n* is the number of IDRs in the Training set and |*SHAP_value_*| measures the importance of each feature for the model decisions.

### Dimensionality reduction analysis of the Training IDRs and human IDRome

To validate whether the IDR-Puncta ML model can be applied to the human IDRome for predicting probability of condensate formation and puncta classification. We performed dimensionality reduction analysis by applying Uniform Manifold Approximation and Projection (UMAP)^64^ using the 25 physicochemical features of the Training IDRs and human IDRome combined. UMAP algorithm aims to preserve both the local and the global data structure. The IDRs from human IDRome with missing feature(s) were excluded from the UMAP analysis. The feature values of the Training IDRs and human IDRome were converted to z-score before applying UMAP. UMAP analysis was performed using the *uwot* package (version: 0.1.12) in R with the parameters, *n_neighbors* = 100, *metric*=“euclidean”, *min_dist* = 1, and *spread* = 5.

### GO enrichment analysis of human IDRome

GO enrichment analysis of the human IDRome was performed using the Database for Annotation, Visualization and Integrated Discovery (DAVID)^65^ Knowledgebase (version: v2024q1) in the web server (david.ncifcrf.gov). For the predicted puncta(+) IDRome, proteins with at least one puncta(+) IDR were used, whereas for the predicted puncta(-) IDRome, proteins with only puncta(-) IDRs were used, and proteins from the full human proteome were used as the background set. The lowest level of GO categories for biological process (GO_BP_ALL), cellular component (GO_CC_ALL) and molecular function (GO_MF_ALL) were used in DAVID. The adjusted p-value (false discovery rate) cutoff of ≤ 0.05 based on Benjamini correction (default in DAVID) and fold enrichment of ≥ 1.5 was used to filter the GO terms. Additionally, GO terms containing ≥ 2,000 proteins from the background set (human IDRome) and GO terms containing < 10 proteins from the target sets [predicted puncta(+) and puncta(-) IDRomes] were removed. The *rrvgo*^66^ package (version: 1.10.0) was used to simplify the redundance of GO terms by grouping similar terms based on their semantic similarity using the “Wang” method with similarity threshold values of 0.88 and 0.94 for the predicted puncta(+) and puncta(-) IDRomes, respectively. “Wang” is a graph-based method implemented in the *rrvgo* R-package, which uses the topology of GO graph structure to compute semantic similarity^67^. This method determines the semantic similarity of two GO terms based on both locations of these terms in the GO graph and their relations with their ancestor terms. Higher thresholds lead to fewer groups of GO terms. For visualization purposes, the combined adjusted p-value of a grouped GO term from semantic similarity was obtained by applying the sum of logs (Fisher’s) method using the *metap* (version: 1.11) R-package. A *log_2_* transformation was used to calculate the average fold enrichment of grouped GO terms.

### Puncta prediction of the IDRs from proteins in membrane-less organelles (MLOs) and membrane contact sites (MCSs)

Protein constituents of MLOs were obtained from the manually curated database of phase-separation related proteins (PhaSepDB; version: 2.1^48^). This database contains 499 human proteins identified by low throughput methods, termed “MLO-lt”, that matched UniProt IDs found in our human proteome database. Of these 499 proteins, 345 were mapped onto our human IDRome database. We identified 199 human proteins from a manually curated database of experimentally supported MCS proteins and complexes (MCSdb^49^) labeled with “low throughput experimental methods”. Two out of the 199 proteins were not found in our human proteome database, and for the remaining 197 proteins, 90 proteins were mapped onto our human IDRome database. Significance of the number of proteins containing puncta(+) or only puncta(-) IDRs in MLO, MCs and hman IDRome data sets were computed from Fisher’s exact test for count data using *fisher.test* from the *stats* (version: 4.2.1) package in R.

## Acknowledgments

We thank Dr. Ines Chen for the critical review of the manuscript and Dr. Steven W. Whitten for providing the AAindex classification. This work is supported by NCI R01 CA246125 (to R.W.K. and M.M.B.), NCI U54 CA243124 (to R.W.K.), Developmental Funds under NCI P30 CA021765 (to R.W.K.) and ALSAC. We are grateful for the support of Core Facilities used in this study by NCI P30 CA021765, including the Cell and Tissue Imaging Center, with technical support from George Campbell, Aaron Taylor, and Aaron Pitre, and the Hartwell Center. This research content is solely the responsibility of the authors and does not necessarily represent the official views of the National Institutes of Health.

## Author Contributions

Conceptualization, SM, ST and RWK; Software, ST, DWB, JB, RS and BL; Investigation, SM, ST, DWB, AHP, CP, JW, W, WTF, SK, BP, RS and BL; Writing - Original Draft, SM, ST and RWK; Writing – Review & Editing, all authors; Supervision, RWK and MMB; Funding acquisition, RWK and MMB

## Corresponding author

Correspondence to Richard W. Kriwacki.

