## Supplementary Figures for "Proteome-wide computational analyses reveal links between protein condensate formation and RNA biology"

**Supplementary Figures and Legends**

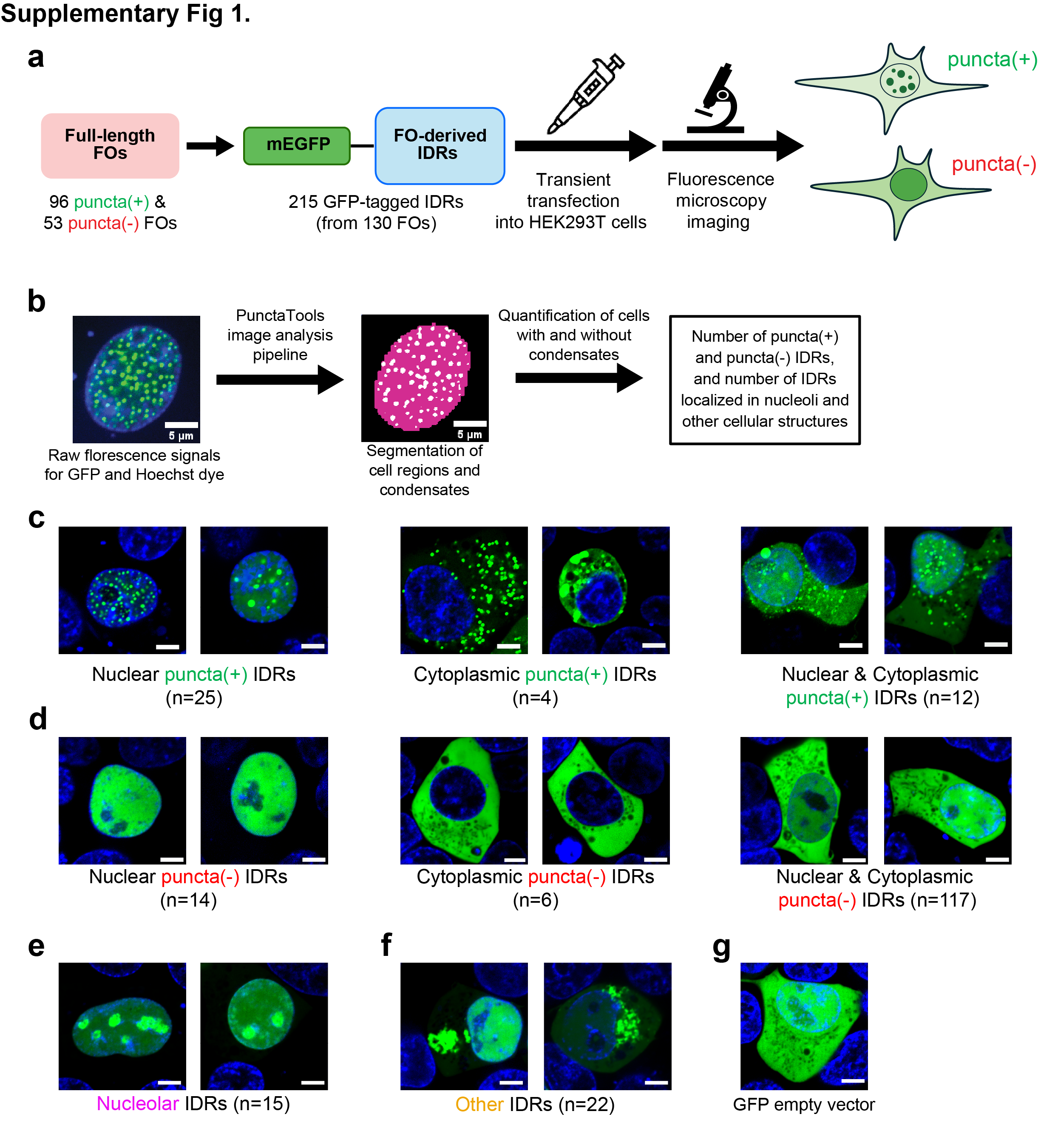

**Supplementary Fig. 1. Representative live cell images of mEGFP-tagged IDRs. (a)** Schematic representation of the IDR imaging workflow. A total of 215 IDRs were analyzed for condensate formation in HEK293T cells, termed the Expressed IDRs. **(b)** Example of puncta segmentation by PunctaTools for quantitative analysis of cells with and without condensates (see Methods). **(c)** Representative confocal microscopy images of live HEK293T cells expressing mEGFP-tagged puncta(+) IDRs localized to the nucleus (left), cytoplasm (middle) or both compartments (right) based upon two biological replicates. **(d)** Representative confocal microscopy images of live HEK293T cells expressing mEGFP-tagged puncta(-) IDRs localized to the nucleus (left), cytoplasm (middle), or both (right). **(e-f)** Representative confocal microscopy images of live HEK293T cells expressing mEGFP-tagged IDRs classified as Nucleolar (e) or Other (f), based upon two biological replicates. The numerical values in (c-f) indicate number of IDRs in different cellular locations (in parentheses). **(g)** Representative confocal microscopy images of live HEK293T cells expressing mEGFP empty vector as a negative control based upon two biological replicates. In all images, the IDR signal (green) is overlayed with the DNA signal (Hoechst dye, blue). All scale bars are 5 μm.

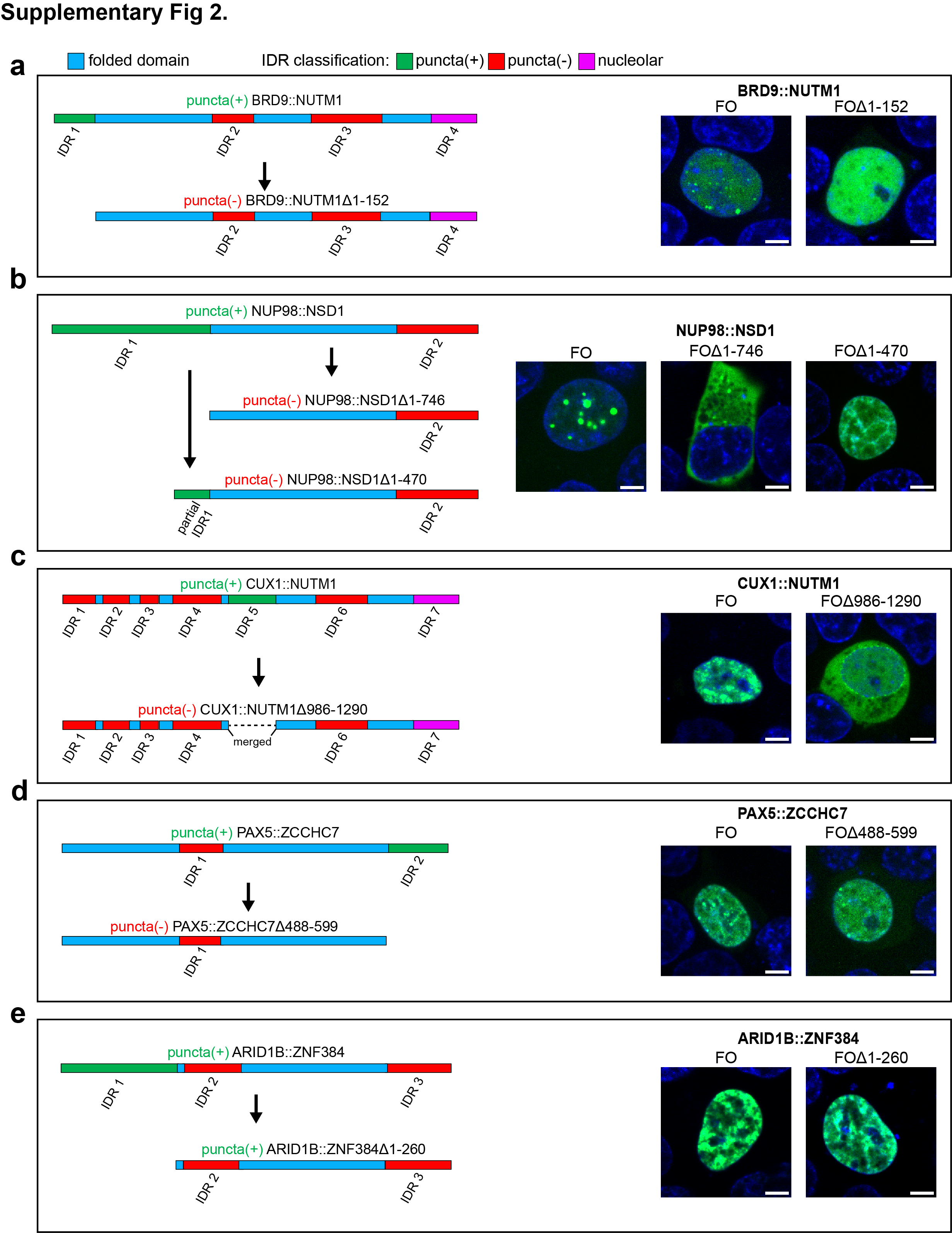

**Supplementary Fig.2. Representative live cell images of mEGFP-tagged FO-IDR deletion mutants.** Schematic showing full or partial deletion of a puncta(+) IDR from full length FOs (left) and their representative confocal microscopy images of live HEK293T cells expressing mEGFP-tagged full-length FO and FO-IDR deletion mutants [puncta(+) IDR deleted FOs], based upon two biological replicates (right) for **(a)** BRD9::NUTM1**,** **(b)** NUP98::NSD1, **(c)** CUX1::NUTM1, **(d)** PAX5::ZCCHC7, and **(e)** ARIDIB::ZNF384. The full-length FOs (BRD9::NUTM1, NUP98::NSD1, CUX1::NUTM1, PAX5::ZCCHC7, and ARIDIB::ZNF384) formed nuclear puncta. Several puncta(+) IDR deletion mutants, including BRD9::NUTM1(Δ1-152), NUP98::NSD1(Δ1-746), CUX1::NUTM1(Δ986-1290), and PAX5::ZCCHC7(Δ448-599), exhibit diffuse localization. The ARIDIB::ZNF384(Δ1-260) mutant exhibited similar condensation behavior to the full-length protein. For NUP98::NSD1, deletion of the entire puncta(+) IDR (residues 1-746, Δ1-746 ) resulted in a diffuse cytoplasmic localization. To investigate this FO further, we generated an additional NUP98::NSD1 deletion mutant where we specifically removed the NUP98 portion (residues 1-470, Δ1-470), which includes the FG repeats required for condensation. This mutant displayed a diffuse nuclear localization as well as nucleolar partitioning. In all images, the protein signal (green) is overlayed with the DNA signal (Hoechst dye, blue). All scale bars are 5 μm.

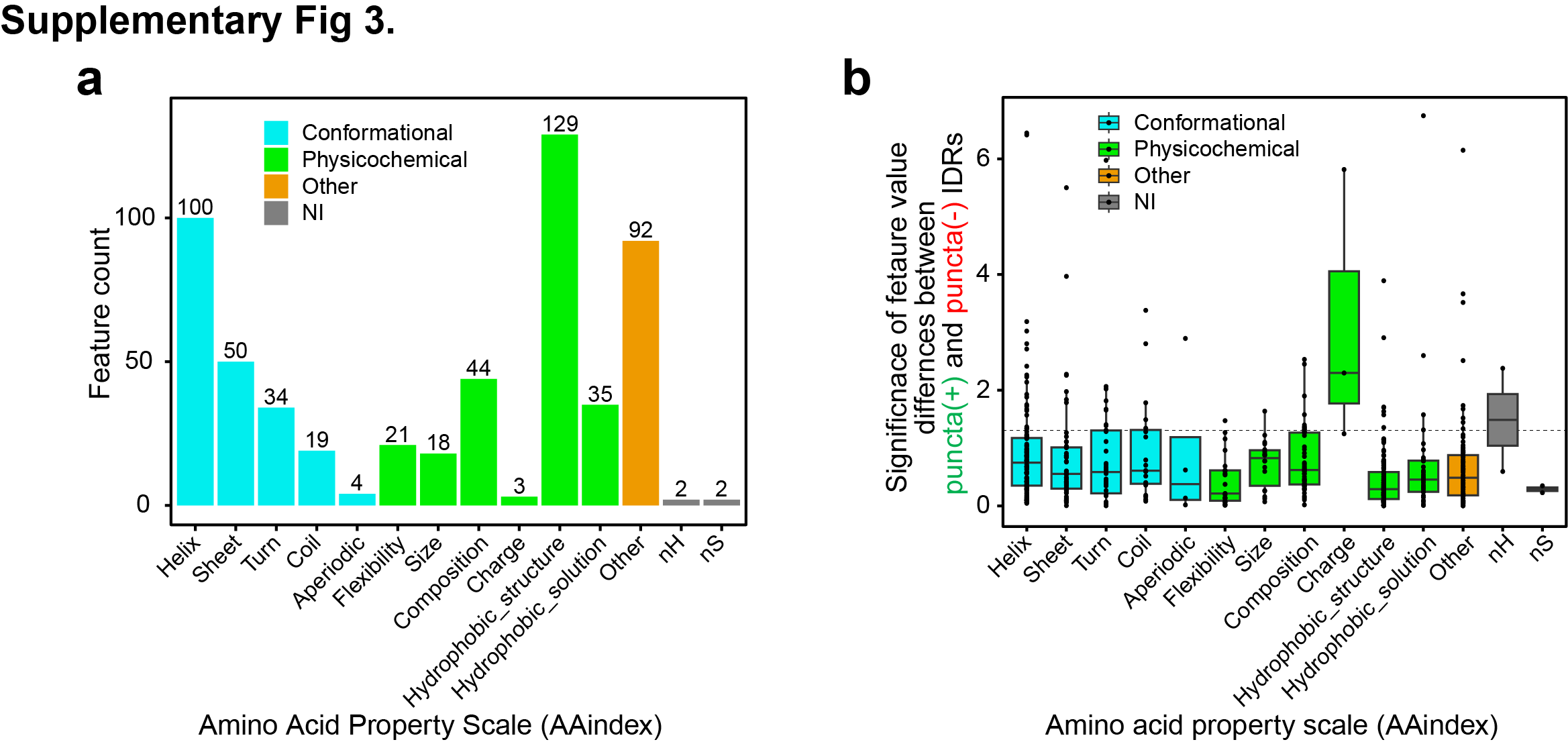

**Supplementary Fig. 3. Classification and significance of the AAindex features. (a)** Classification of the 553 AAindex features without missing values into 12 classes that are broadly divided into four categories, Conformational (cyan), Physicochemical (green), Other (orange) and NI (features not included in our study; grey). **(b)** Significance of differences in the average values of 553 AAIndex features for puncta(+) and puncta(-) IDRs, reported as -log_10_(p-value) determined using the two-sided Welch’s *t*-test with no adjustments for multiple comparisons. The dotted horizontal line indicates p-value=0.05. The features in two classes “nH” (non-helical) and “nS” (non-sheet) were not included in this study following work of Ibrahim *et. al*^25^.

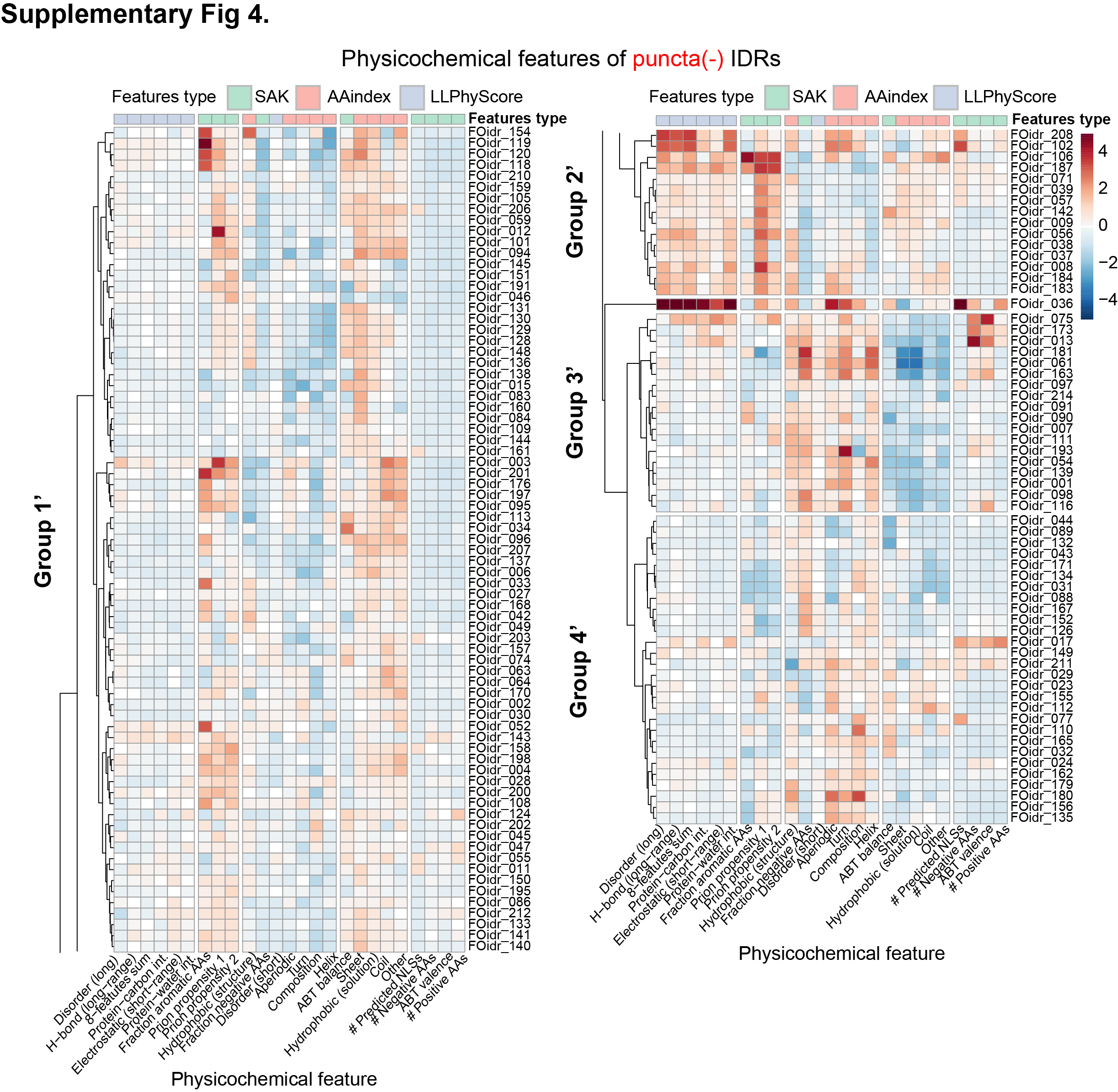

**Supplementary Fig. 4. Heatmap groups of the puncta(-) Expressed IDRs.** Results of two-dimensional (2D) hierarchical clustering of the 137 puncta(-) IDRs based on the 25 most discriminatory physicochemical features as z-scores (columns) with respect to the human IDRome (see Methods) into four groups (Groups 1’–4’). The top row represents feature types. IDR feature values are color-coded in the rows, with IDR names given on the right. The names of the physicochemical features are given at the bottom. Note that a single IDR (FOidr_036) forming an independent cluster between Group 2’ and Group 3’ was not considered for analysis.

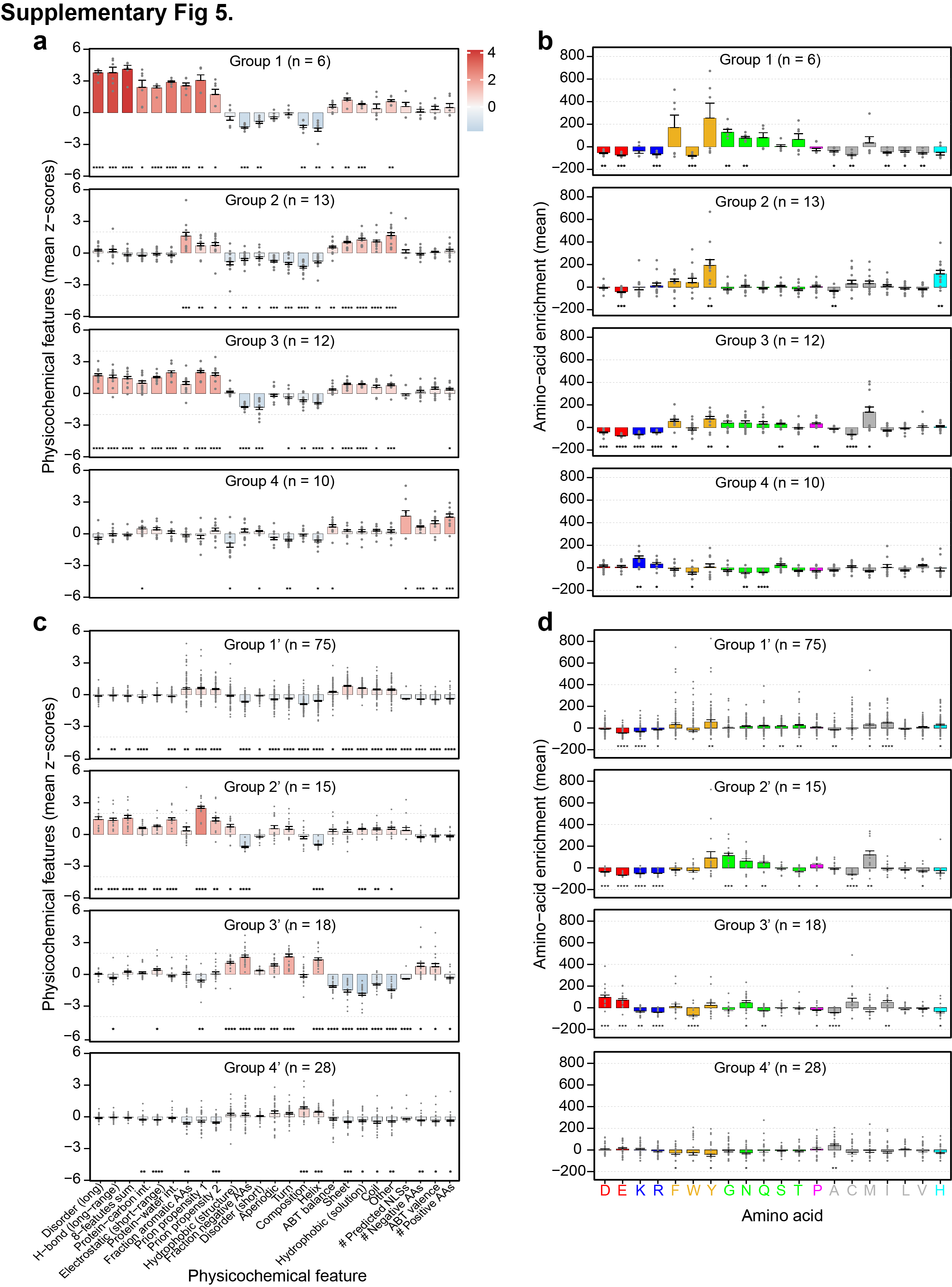

**Supplementary Fig. 5. Average physicochemical feature values and amino acid enrichments for the puncta(+) and puncta(-)IDRs on the basis of the feature groups. (a)** Quantification of the mean enrichment or depletion values for the 25 physicochemical features for Groups 1–4 for 41 puncta(+) Expressed IDRs (see **Fig. 2c**). Values are reported as mean z-scores ± standard error and normalized with respect to the human IDRome. **(b)** Quantification of the mean amino acid enrichment or depletion ± standard error for IDR sequences relative to the human IDRome in Groups 1–4 for 41 puncta(+) Expressed IDRs. **(c)** Quantification of the mean enrichment or depletion values for the 25 physicochemical features for Groups 1’–4’ for 137 punta(-) Expressed IDRs (see **Supplementary Fig. 3**). Values are reported as mean z-scores ± standard error and normalized to the human IDRome. **(d)** Quantification of the mean amino acid enrichment or depletion ± standard error for IDR sequences in Groups 1’–4’ for 137 punta(-) Expressed IDRs relative to the human IDRome. In (a-d), the value n in parentheses indicates the number of IDRs in the corresponding feature group.

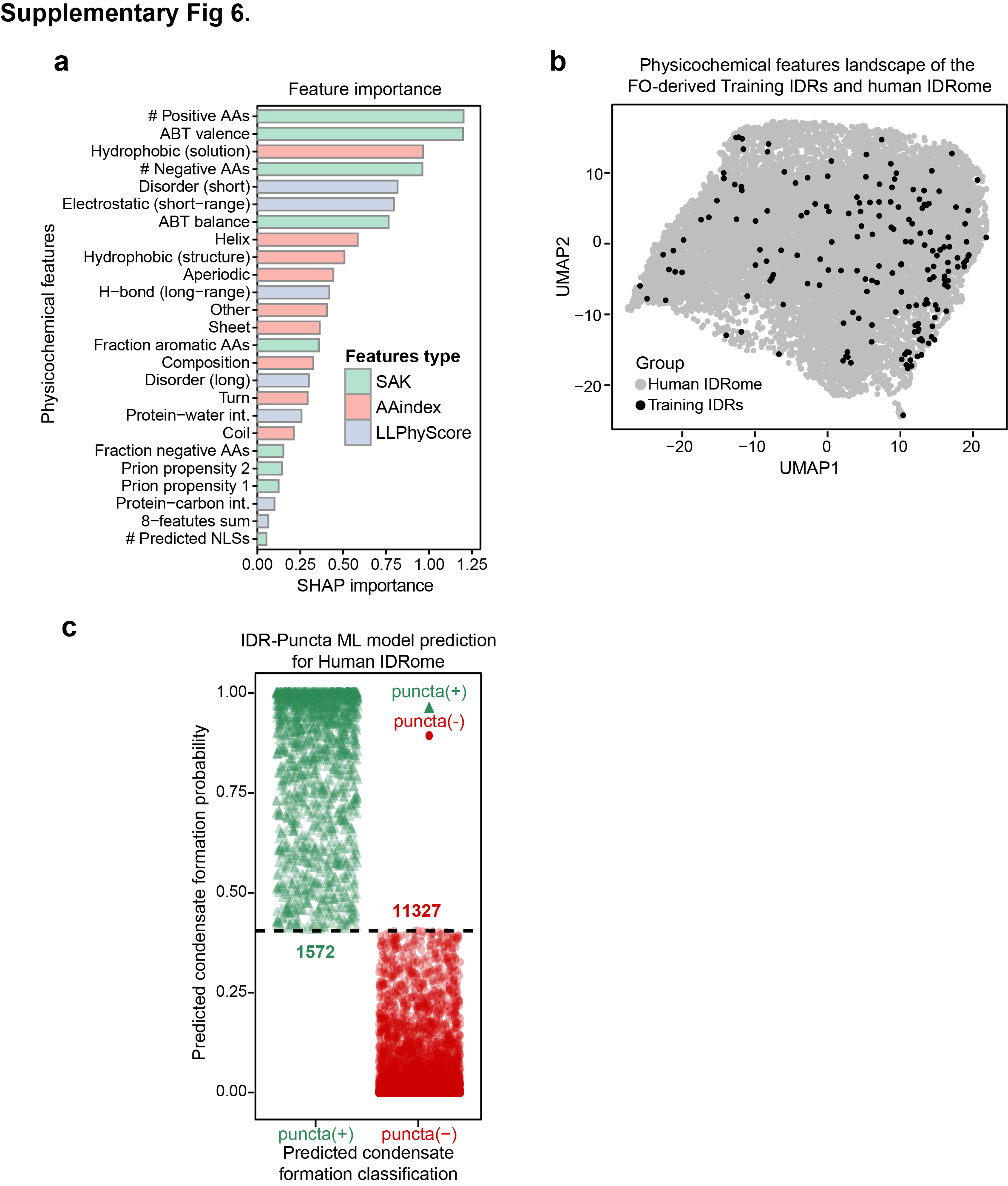

**Supplementary Fig. 6. Feature importance, performance metrics and condensation behavior prediction from the machine learning model. (a)** SHapley Additive exPlanations (SHAP) analysis for feature importance for predictions made using the base model gradient boosting machine (GBM) from the IDR-Puncta ML model (see Methods) based on analysis of the 178 Training IDRs [41 puncta(+) and 137 puncta(-) Expressed IDRs] colored by feature types (SAK, cyan; AAindex, red; LLPhyScore, blue). The features are ranked by the magnitude of their relative SHAP contributions to predictions, with those with the largest contributions toward the top. **(b)** Uniform Manifold Approximation and Projection (UMAP) analysis of the Training IDRs and human IDRome using the 25 physicochemical features as z-score (see Methods). On the reduced two-demensional (2D) projection of UMAP plot, 178 Training IDRs and 12,276 IDRs from human IDRome [with non-missing feature (s)] are represented in black and grey dots, respectively. **(c)** Results of predicted condensation behavior using the IDR-Puncta ML model for all IDRs in human proteome termed human IDRome (12,899 unique IDRs, in total). Predicted condensate formation probability from the IDR-Puncta ML model is along the y-axis. The predicted puncta(+) IDRs are depicted with green triangles, while the predicted puncta(-) IDRs are depicted with red circles. The IDR-Puncta ML model threshold for puncta(+) classification is a probability value greater than or equal to 0.4, and otherwise classified as puncta(-) (indicated by the dashed horizontal line).

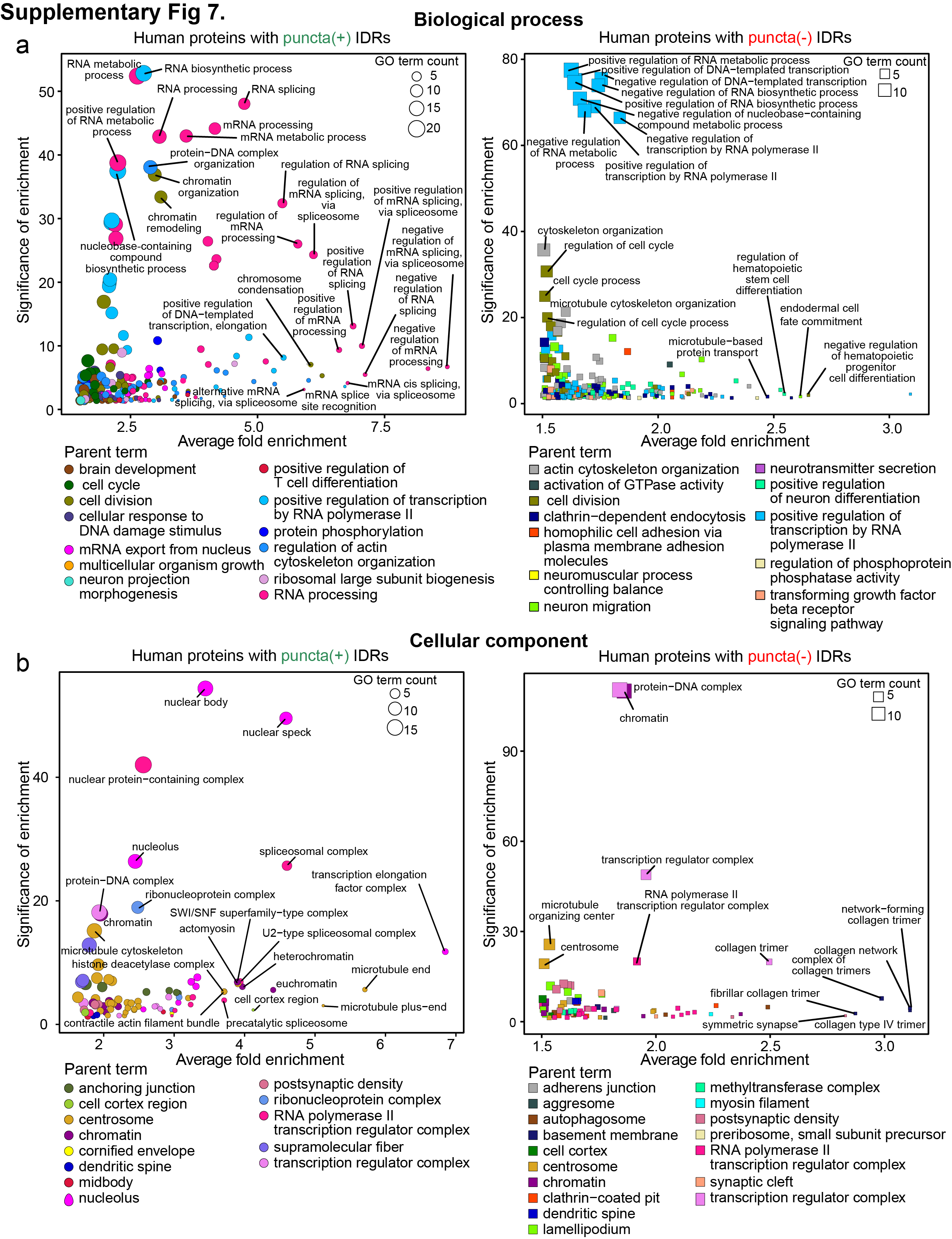

**Supplementary Fig. 7. GO term enrichment analysis of human IDRome. (a)** GO terms related to biological processes of proteins with predicted puncta(+) IDRs (left) and proteins with only predicted puncta(-) IDRs (right). **(b)** GO terms related to cellular component of proteins with predicted puncta(+) IDRs (left) and proteins with only predicted puncta(-) IDRs (right). In (a) and (b) GO enrichment analyses are summarized as scatter plots [x-axis, fold enrichment]. The adjusted p-value (cutoff ≤ 0.05 using *Benjamini correction*) is along the y-axis. Each symbol represents a GO term, and the colors indicate the grouping of the GO terms based on their semantic similarity in the legend below the plots (see Methods). The size of each GO term symbol indicates the number of proteins from the target list in that GO term.

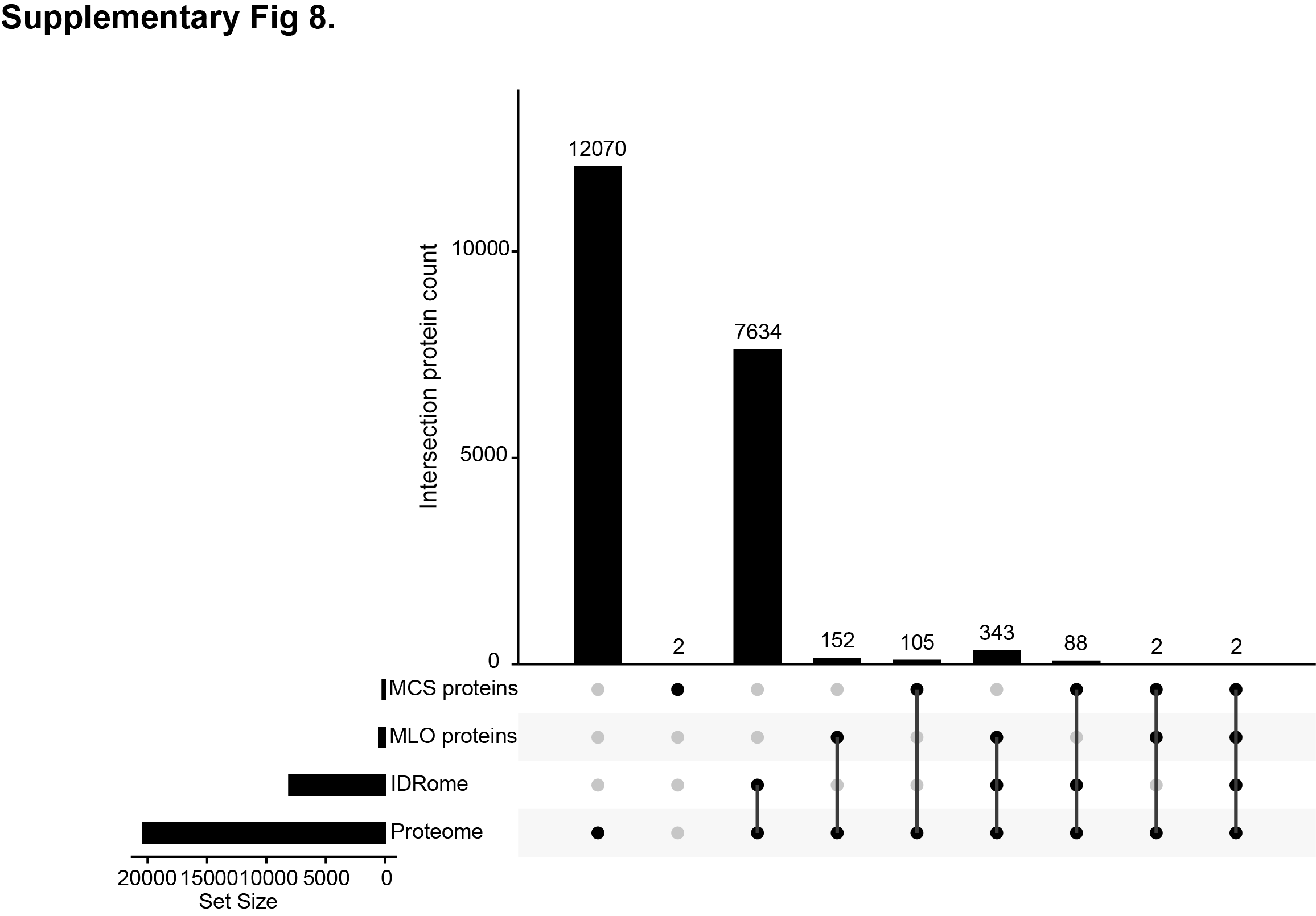

**Supplementary Fig. 8. Analysis of overlap between human proteome, proteins with IDRs, MLO proteins and MSC proteins.** UpSet plot showing intersection of the human protein sets used in our analyses, including human MLO proteins (high confidence), MCS proteins (high confidence), entire proteome, and proteins with IDRs (IDRome).

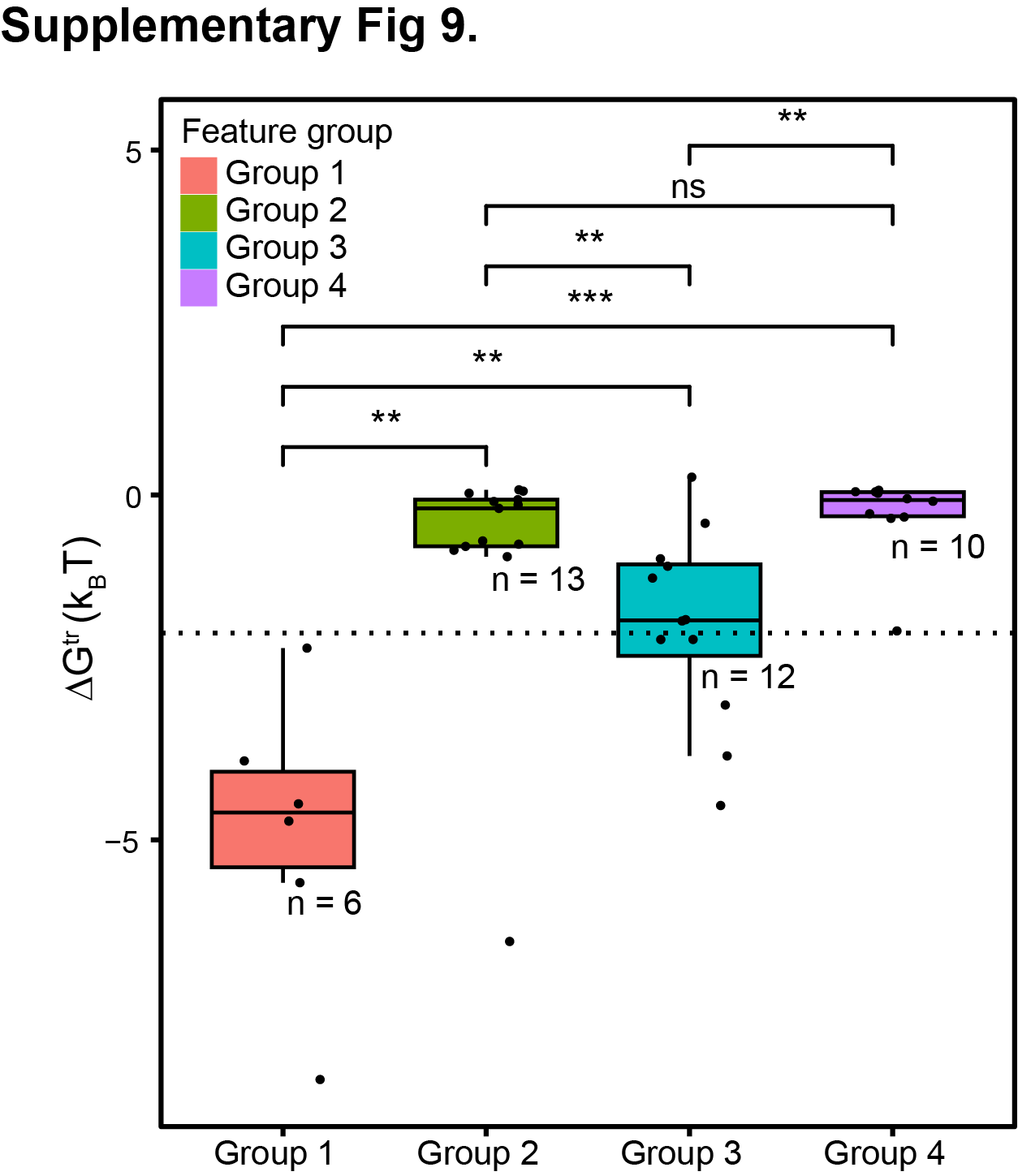

**Supplementary Fig. 9. Prediction of free energy for phase separation of the 41 puncta(+) Expressed IDRs from the four heatmap groups.** Quantification of predicted Gibbs free energy of transfer (ΔG^tr^; units of k_B_T) for the Expressed 41 puncta(+) IDRs as calculated using a ML model derived from coarse-grained molecular dynamics simulations (by von Bulow, et. al.). ΔG^tr^ quantifies the thermodynamic favorability of condensate formation by polypeptide chains. Violin plots displaying the distribution of ΔG^tr^ values for IDRs in each of the four puncta(+) feature groups (see **Fig. 3c**). Statistical significance of differences between the groups was determined using a two-sided Welch’s *t*-test (no adjustments were made for multiple comparisons) and indicated by asterisks (*p < 0.05, **p < 0.01, ***p < 0.001, ns = not significant). The dotted line represents a ΔG^tr^ value of -2 k_B_T.
